# Protein Translation Dysregulation and Immune Cell Evasion Define Metastatic Clones in HPV-related Cancer of the Oropharynx

**DOI:** 10.1101/2024.04.15.589624

**Authors:** Venessa T Chin, Walter Muskovic, Rachael A McCloy, Drew R Neavin, Jose Alquicira-Hernandez, Himanshi Arora, Anne Senabouth, Patricia Keith, Ellie Spenceley, Angela Murphy, Dominik Kaczorowski, Peter Floros, Peter Earls, Brett Leavers, Julia Crawford, Richard Gallagher, Joseph E. Powell

**Affiliations:** Translational Genomics, Garvan Institute of Medical Research, Darlinghurst, Sydney, NSW, Australia; The Kinghorn Cancer Centre, St Vincent’s Hospital Sydney, Darlinghurst, Sydney, NSW, Australia; Faculty of Medicine, University of New South Wales, Kensington, NSW, Australia; Faculty of Medicine, University of Notre Dame, Sydney, NSW, Australia; Cellular Genomics Futures Institute, University of New South Wales, Kensington, Sydney, NSW, Australia

**Keywords:** Oropharyngeal cancer, HPV, clonal evolution, metastasis, tumour microenvironment, translational relief

## Abstract

Head and neck cancers, representing the seventh most common malignancy globally, have seen a shift in causative factors from traditional smoking and alcohol use to human papillomavirus (HPV) infection, now accounting for up to 80% of oropharyngeal cancers. We identify the cellular and clonal mechanisms underlying immune avoidance and metastasis by analysing single-cell and spatial genomic data from primary and metastatic cancers. We first map the clonal evolution of malignant cells based on the accumulation of mutations. We identify metastasising clones based on mutational similarity scores between cells in the primary and lymph node metastasis. Genomic analysis of metastasising and non-metastasising clones identified virally mediated protein translation relief (*P*=4.24x10^-24^) pathway underlying metastatic expansion. We show that in metastatic clones, this process is driven through upregulation of transition-initiating factors, *EIF4E* (*P*=1.5x10^-13^) and *EIFG1* (*P*<2.22x10^-16^), and suppression of regulatory kinases *EIF4EBP1* (*P*=2.1x10), *EIF2AK2* (*P*<2.22x10^-16^), and *EIF2S1* (*P*<2.22x10^-16^). We subsequently identify that metastatic clones have a corresponding downregulation of the *JAK/STAT* pathway and immunoproteasome genes *PSMB8* (*P*<2.22x10^-16^) and *PSMB9* (*P*<2.22x10^-16^), suggesting these clones escape immune surveillance through decreased *INF* inflammatory response and antigen presentation. We validate these results using spatial RNA-seq data, where metastatic cancer clones show decreased cell-to-cell interactions with CD4 T-effector memory cells (CD4_TEM_) (*P*=0.0077), CD8 T-exhausted cells (CD8Ex) (*P*=0.0191), and innate lymphoid cells (ILC) (*P*=0.04). Finally, we demonstrate that the upregulation of cap-independent translational drives cell proliferation in metastatic clones through the expression of translation initiation factors (*EIF4G1: P*<2.22x10^-16^). Our results provide evidence of the mechanisms by which virally induced cancer clones lead to advanced disease and poor prognosis in patients.

## Introduction

Cancers of the head and neck comprise a diverse range of cancers, which represent the 7^th^ most common malignancy worldwide, with approximately 900,000 new cases and 450,000 deaths per year^1^. Historically, cancers in the oropharynx region were commonly associated with smoking and alcohol use. However, infection with human papillomavirus (HPV) is becoming an increasingly common cause and is now responsible for up to 80% of oropharyngeal cancers^2^. The HPV-positive oropharyngeal cancer (HPV+) patient population differs from HPV-negative (HPV-), with earlier age of onset, smaller primary tumours, and larger lymph node burden but improved survival outcomes^3–5^. The cellular mechanisms underlying these clinical differences in HPV+ and HPV-patients are poorly understood^6,7^, resulting in missed opportunities to understand the defining differences in cancer cell behaviour, metastasis and treatment response. Despite aggressive therapies, up to 25% of patients with potentially curable HPV+ oropharyngeal cancer experience disease relapse, after which most patients die within two years^8^. To reduce the risk of relapse, adjuvant radiation given with or without chemotherapy can be offered. However, identifying patients who will benefit the most from these toxic therapies with high-risk disease remains difficult without accurate prognostic biomarkers^9^.

Cancer cells are genetically heterogeneous, leading to cancer phenotypes that are transcriptionally and functionally distinct from one another^10^. Primary tumours typically exhibit greater clonal diversity than metastases^11^. This observation is an emergent property because metastatic processes are partly driven by cells acquiring genetic mutations that impact their phenotypes through clonal selection within their microenvironmental context ^12–15^. Identifying the phenotypic characteristics and molecular mechanisms of clones prone to metastasis not only offers a chance to identify biomarkers indicating a poor prognosis but also holds the potential for uncovering novel drug targets.

In HPV-oropharyngeal cancer, the epithelial-to-mesenchymal transition differentiates between primary and metastatic cells^16^. Moreover, *AXL*, known for its role in cell migration, EMT, invasion, and proliferation^17^, along with *AURKB*, a pivotal regulator of mitosis^18^, have both been recognised as significant contributors to the development of invasive metastatic cell phenotypes in HPV-oropharyngeal cancers.

In HPV+ oropharyngeal cancers, there is limited understanding of the cellular pathways underlying metastasis, which are expected to differ from those in HPV-cancers given the unique clinical and aetiological factors. The inability to clear HPV infections stems from complex mechanisms involving the suppression of the host cell’s antiviral response (involving *NFKB*, *STING*, and *IFN*), the sustained viral replication facilitated by the retinoblastoma tumour suppressor (*RB*), and evasion of immune surveillance by inhibiting apoptosis via *TP53*^19^.

To sustain viral replication, HPV employs various mechanisms to manipulate the host cell. Specifically, viral proteins E6 and E7 incapacitate key tumour suppressor regulators like TP53^20^ and RB^21^, disrupting apoptosis and cell cycle regulation. Furthermore, E6 directly interferes with DNA damage repair processes^22^. While these strategies effectively support viral replication, they render the host cell incapable of managing defective DNA damage repair, leading to a gradual increase in genomic instability over time^22^. Ultimately, this increasing genomic instability culminates in malignant transformation^23^. Simultaneously, the virus employs multiple mechanisms to evade the host’s immune responses. This immune evasion enables infected cells to remain undetected and allows for the tolerance of emerging pre-malignant and malignant lesions. Therefore, many mechanisms that maintain viral infection may also contribute to cancer cell evasion of anti-tumour responses, an area that has not been comprehensively explored.

Here, we present an analysis of primary and regionally metastatic cancers in patients with HPV+ oropharyngeal cancer who have undergone curative surgical resection. Using single-cell techniques, we use copy number variation (CNV) to define cancer clones and study clonal population phenotypes with regard to metastasis and microenvironmental dynamics. We identify virally mediated protein translation relief and dysregulation of immunoproteasomal and IFN pathways, accompanied by a shift towards cap-independent translation underlying metastatic clonal selection. Furthermore, our study validates immune cell evasion by sequencing matched tumour samples *in situ*.

## RESULTS

### Evidence of clonal selection favouring phenotypic predisposition to metastasis

It has long been recognised that the genomic profiles of cancer clones influence cellular phenotypes, impacting clinical outcomes^24^. Using haplotypes and single-cell RNA-sequence (scRNA) data, we called CNVs from matched primary and regional lymph node metastases from four HPV+ patients (**Figure 1A**, **Figure 1B, Supplementary Table 1**)^25^. An overview of the CNV distribution across all individual cells highlights various genomic alterations indicative of the genomic instability typically observed in cancers (**Figure 2**). We did not observe a consistent enrichment of oropharyngeal cancer oncogenes identified by the cancer genome atlas (TCGA)^26^ in the CNV regions (*PIK3CA, TRAF3, E2F1*) (**Figure 2**). A possible explanation is that TCGA data comprises HPV+ and HPV-cancer samples. Nevertheless, the data illustrates the heterogeneous nature of HPV+ cancer across patients. To understand clonal dynamics, subclonal phylogenies were generated by *Numbat*^27^ by aggregating data from single cells into a pseudobulk profile, capturing the evolutionary relationships between cells. From 14,157 single-cell CNV profiles, three to seven distinct clusters per patient were identified (**Figure 3A**). Each cluster, suggesting a unique cancer clone, exhibited a characteristic CNV signature. By constructing a phylogenetic tree based on shared CNV events^25^, we identified the ancestral clones and genomic evolution of clones by acquiring new somatic events. As expected, the clonal evolution of cancers was unique for each patient and is summarised in detail in **Figure 3**. Using patient four as an example, we observe the development of clone two arising from the original clone (ancestral) through deletions at 3p and 15p. Subsequent deletions at 10q and 11q and a loss of heterozygosity (LOH) at 20q led to clone three, followed by a divergent event into clones four and six through a new LOH at four (clone six) and amplifications at 3q and 5q plus a deletion at 7q (clone four). We observe a final clonal evolution event, with clone five arising from clone four via somatic mutations at 4p^+^, 8q^-^, 19^+^ and 20p^+^ (**Figure 3**). Similar patterns of clonal evolutionary events are observed across patients.

**Figure 1:**
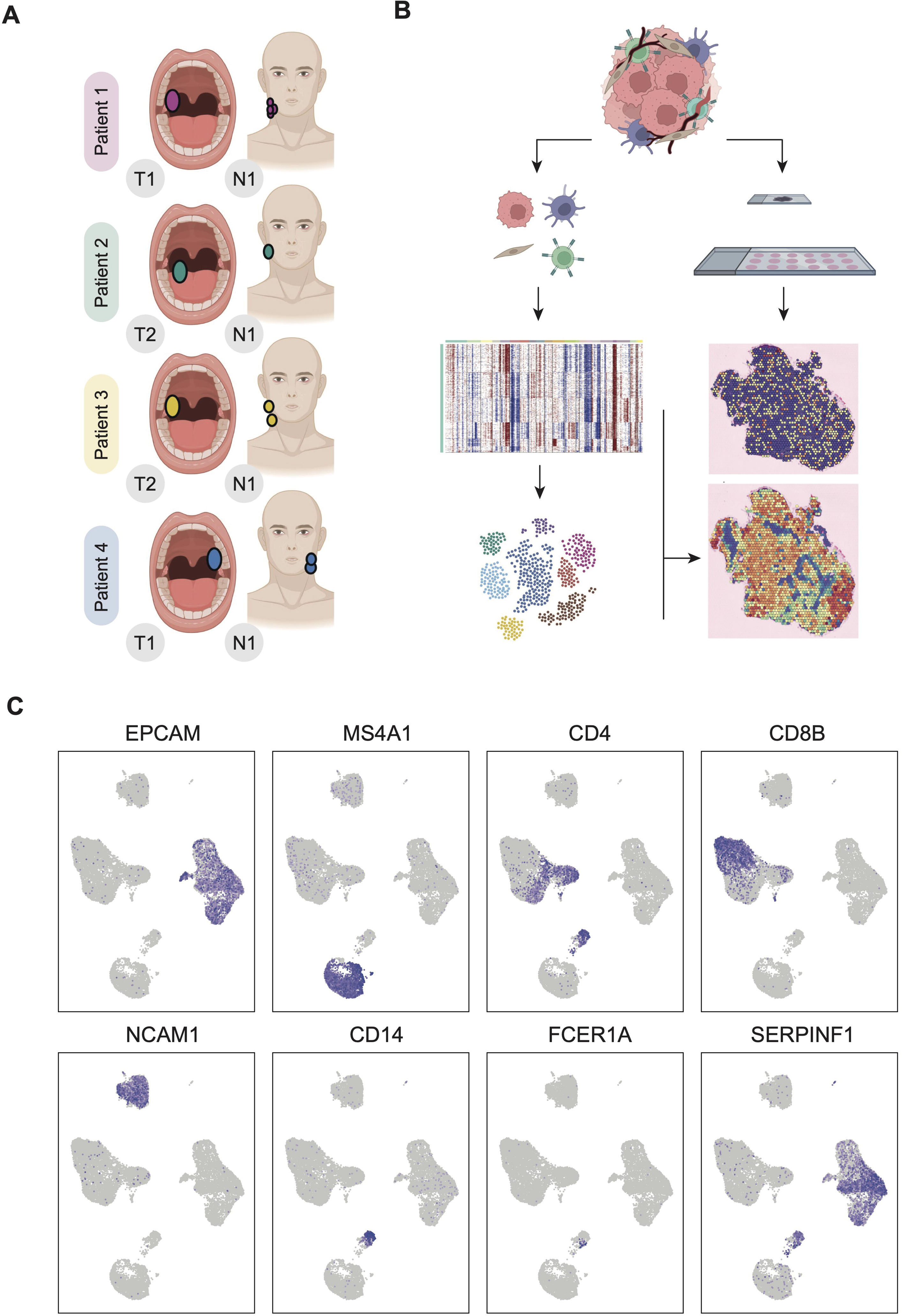
Study design and cell annotation. A. Description of the patient samples. The primary sites of the oropharyngeal of either the tonsil or base of the tongue, nodal stations affected by cancer as per the pathological staging. B. Study schematic. In parallel, the samples were processed for scRNA-seq and spatial RNA sequencing using the Visium platform. Annotated single cells were used to deconvolute the spatial data. Clone delineation was performed based on single-cell CNV calls. C. Canonical markers demonstrate cancer cells, b-cells, t-cells, NK cells and monocytes.

**Figure 2.**
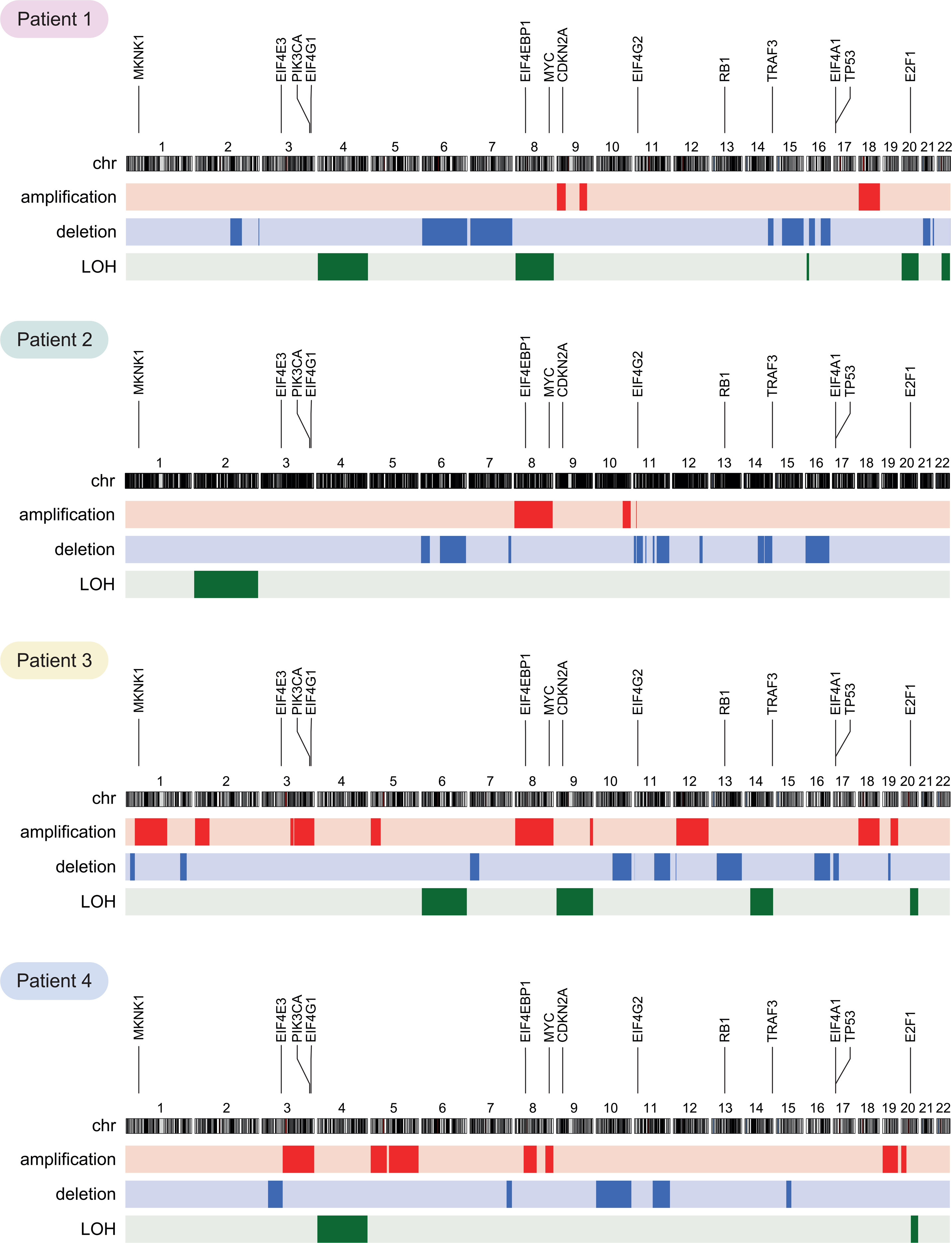
Karyotype plots. For each patient, the karyotype plots, including the primary and the LN, are shown—amplification in red, deletion in blue and LOH in green. The location of key genes are shown at the top. Chromosome 8 was shown to be altered in all patients. Locations of genes of interest are listed across the top. HPV infection and cancer (*RB1, TP53, MYC*), genes identified in the ICGC cohort (*E2F1, TRAF3, PIK3CA*) and translation initiation (*EIF4E3, EIF4G1, EIF4EBP1, EIF4G2, EIF4A1, MKNK1*)

**Figure 3:**
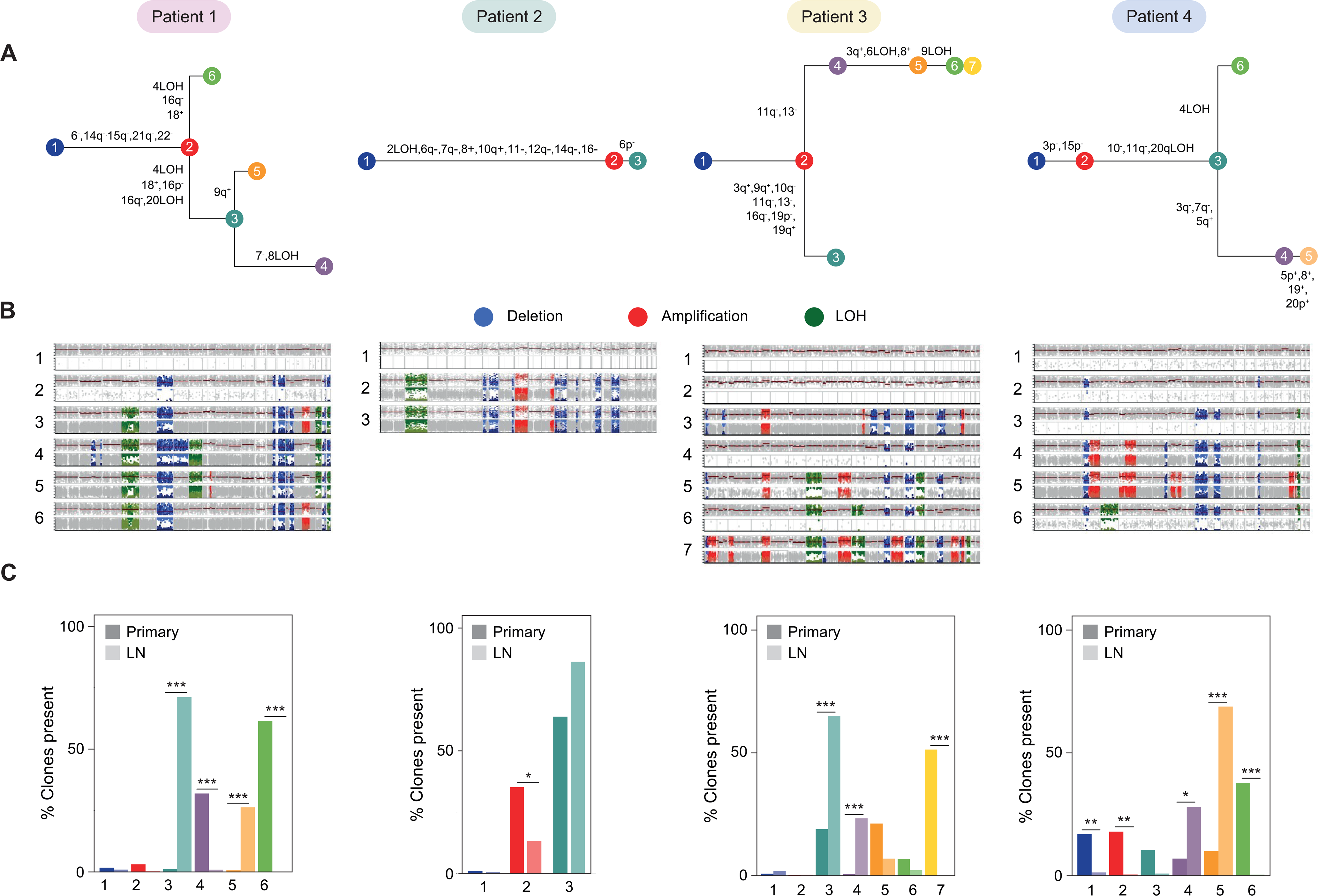
Selection leading to clonal expansion in metastatic lymph nodes. A. Clonal phylogenies are shown for each patient. B. CNV events underlying the formation of expanding clones. Each row displays the CNV profile for a clone. Deletion, amplification and loss of heterozygosity (LOH) events are shown in blue, red, and green, respectively. C. Percentage representation of each clone in the primary (dark colours) versus the LN (light colours). Significant changes in the percentage representation of a clone between the primary and LN were estimated using a t-test

We next sought to understand the clonal diversity in primary and lymph node sites and identify cancer clones that show significant expansions into the metastatic lymph node sites. As shown in multiple cancers^11^, the clonal diversity was greater in the primary site compared to the lymph metastasis (**Figure 3C**). This difference in diversity can be due to several factors, such as selection pressures on cancer clones, where only a subset of cells from the primary tumour possess the necessary genomic and phenotypic characteristics required to metastasise; tumour microenvironment and adaptation, where differences in immune evasion in the primary and metastatic site favour growth and expansion of specific clones. We observed significant differences in the clonal proportions between the primary and lymph node sites, demonstrating clonal expansion into the metastatic site. This enabled us to identify genomic factors underlying metastatic behaviour. The results revealed a clear pattern, with one or two clones displaying substantial expansion from rare populations in the primary to dominant clones in the metastatic lymph node (**Figure 3**). We refer to these clones as expanding or metastatic clones. Correspondingly, clones common in the primary site are frequently absent or observed in very low frequencies in the lymph node site, suggesting an absence of metastatic phenotypes. We subsequently analysed the genomic differences between these clones to identify pathways and microenvironmental contexts that contribute to the metastatic phenotype.

### Virally-induced dysregulation of protein translation is a defining characteristic in metastasising cancer clones

In most cases, cancer mortality is primarily attributed to metastatic disease^28^, due to the emergence of cancer cell clones with aggressive phenotypes through a selective process. To identify the genomic phenotypes of expanding clones, we overlapped the clonal data with cell clustering to delineate the expanding and non-expanding cancer cells **(Figure 4A)** and identify genes differentially expressed. We identified significantly differentially expressed between the expanding and non-expanding clones (**Supplementary Table 2**). We performed pathway enrichment on this set of genes (**Figure 4B**). Our results identify three broad cellular pathway groups represented by genes whose expression levels significantly differ between metastatic expanding clones and non-expanding clones: cell cycle, immune regulation, and protein translation (**Figure 4B** and **Supplementary Table 3**). In the top ten enriched pathways for each patient comparison, protein translation pathways show consistent downregulation in the expanding compared to non-expanding clones (**Figure 4, Supplementary Figures 1-6**). For example, the pathway “Formation of a pool of free 40S subunits” is enriched in comparison A for patient 1 with 59 differentially expressed genes from this pathway (*P*-value for enrichment = 4.24x10^-23^). **Supplementary Figure 7** shows that the genes found in this pathway in comparison 1A consist of protein translation initiating factors and ribosomal subunit protein genes. Further analysis (**Supplementary Figure 1**) demonstrates that many genes identified in comparison 1A within this pathway are downregulated, like *RPL35A* (Log_2_(FC) = -0.89, P = 9x10^-251^) and *RPS26* (Log_2_(FC) = -0.59, *P* = 1.45 x 10^-107^), indicating that expanding clones are downregulating protein translation.

**Figure 4.**
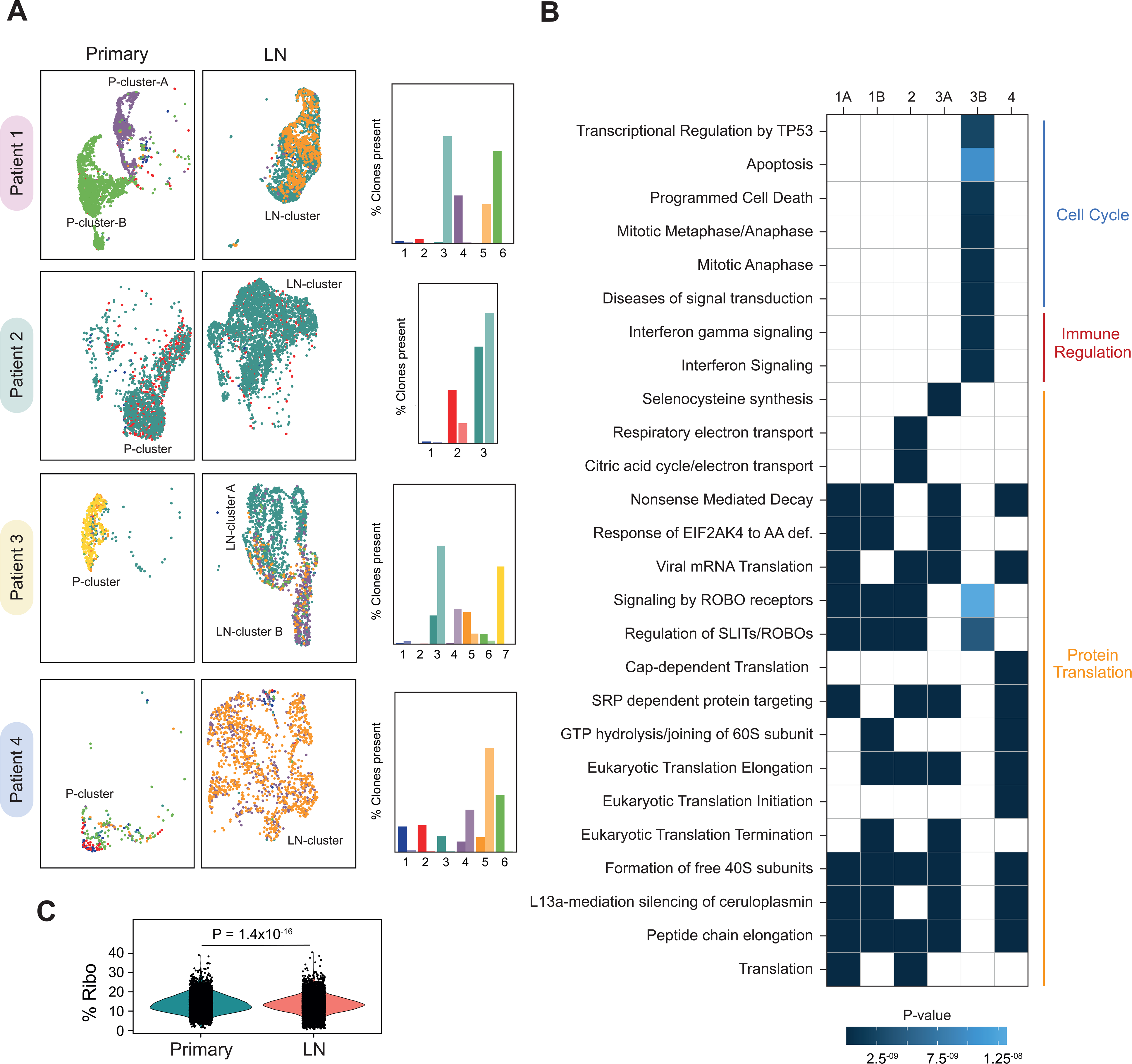
Downregulation of protein translation pathways in expanding metastatic clones. A. The primary and LN cancer cells were clustered into transcriptionally similar groups, with the clones overlayed in their colours shown in the bar graphs. In most cases, the clones formed transcriptionally distinct clusters, which formed the comparisons for pathway enrichment analyses. Multiple comparisons have been done in patients 1 and 3 using these distinct clusters. B. Reactome pathways, which were significantly (study-wide p<0.05) enriched, are shown. The colour relates to the p-value of enrichment. These are grouped into three distinct pathways of cell cycle, immune regulation and protein translation. C. Percentage of ribosomal protein gene expression in the expanding versus non-expanding clones. There is less expression in the expanding clones. A t-test shows that this difference is highly statistically significant. P-cluster = primary cluster. LN-cluster = lymph node cluster.

The consistent downregulation of protein translational machinery in expanding clones is accompanied by a small but significant decrease in the expression levels of ribosomal protein-encoding genes in the expanding clones (*P*=1.4x10^-6^) (**Figure 4C**). The presence of 18 enriched protein pathways, as revealed in **Figure 4B**, all characterised by an abundance of protein translation initiation factors and ribosomal protein genes, collectively underscores the pivotal role of protein translation downregulation as a defining phenotypic factor in expanding clones.

To understand the relationship between viral infection and dysregulation of protein translation in metastatic expanding clones, we tested for the relative expression levels of translation-initiating factors and regulatory kinases associated with virally activated translational relief, a process by which virally infected cells escape normal anti-viral mechanisms^29,30^. Under normal circumstances, a cell will downregulate protein translation and upregulate apoptosis once a virus is detected. We identified significant upregulation of two transition-initiating factors, *EIF4E* (*P*=1.5x10^-13^) and *EIFG1* (*P*<2.22x10^-16^), in expanded cancer clones of the lymph node compared with non-expanding clones and corresponding significant downregulation of regulatory kinases *EIF4EBP1* (*P*=2.1x10), *EIF2AK2* (*P*<2.22x10^-16^), and *EIF2S1* (*P*<2.22x10^-16^) (**Figure 5**, **Supplementary Table 4**). Further evidence of translational relief is seen with upregulation of the stress kinases *PPP1R15A* (*P*<2.22x10^-16^), *PPP1CA* (*P*=7.9x10^-6^) and *PPP1CB* (*P*<2.22x10^-16^) and downregulation of apoptosis-associated genes *TP53* (*P*<2.22x10^-16^), *FAS* (*P*=2.9x10^-8^) and *BAX* (*P*=1.3x10^-9^) (**Figure 5**, **Supplementary Table 4**). Dysregulation of protein translation has previously been described in both viral infection^29,31,32^ and cancer^33^. After an acute viral infection, the host cell attempts to reduce virus replication through a process called “host shut-off”^34,35^, where ribosomal proteins involved in mRNA translation are downregulated and apoptosis is triggered. In normal circumstances, this process limits viral replication and induces apoptosis in infected cells, thereby containing the infection. However, in persistent viral infection like HPV, these mechanisms fail to eradicate the virus. Our results provide strong evidence of virally induced translational relief in expanding clones, where host cells’ shut-off mechanisms are normalised using stress kinases to keep EIF2AK in its dephosphorylated state. This maintains protein translational machinery and viral replication and turns off apoptotic pathways.

**Figure 5.**
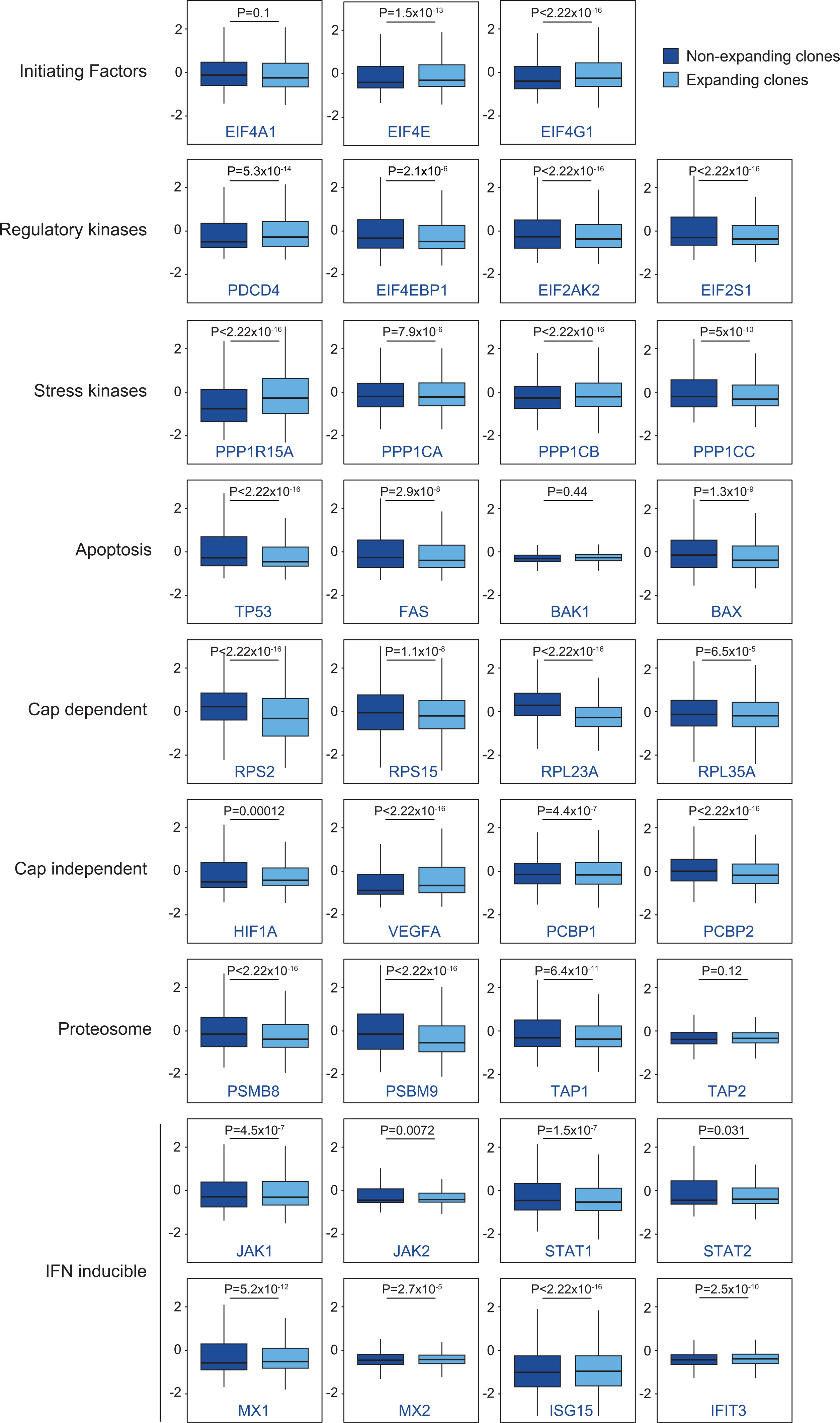
Gene expression differences between metastasising expanding clones and non-expanding clones. Genes are grouped into eight key pathways: Protein translation initiating factors, protein translation regulatory kinases, protein translation stress kinases, apoptosis, cap-dependent translation, cap-independent translation, Proteasome, IFN-inducible genes. The non-expanding cancer clone’s gene expression is in the dark blue and expanding clone’s expression in the light blue.

### Virally-induced immunosuppression through interferon and immunoproteasome pathways

In HPV+ cervical cancer, viral infection triggers significant alterations in the microenvironment^36^. During persistent infection, there is a notable decline in the presence of cytotoxic CD8+ and helper CD4+ T-cells within the epithelial tissue. In the precancerous stage, cytotoxic T-cell populations maintain an immunosuppressed profile, characterised by increased regulatory T-cells. Invasive cancers demonstrate a high immunogenic state marked by elevated levels of cytotoxic and helper T-cells and occasionally include a surge in regulatory T-cell numbers. Notably, abundant regulatory T-cells are associated with a more unfavourable prognosis^36^. Viral infection generally results in immune evasion via suppression of IFN-stimulated gene transcription^32^ and, in HPV-infection, downregulation of the immunoproteasome^30^. Silencing the IFN regulatory pathway has also been demonstrated to be a feature of metastatic cancer cells^37^. We next sought to understand the consequences of HPV infection on immune evasion and its relationship to clonal expansion. Comparing the expression levels of IFN-inducible genes, we observed a downregulation of the IFN regulatory pathway in expanding metastatic cancer clones compared to non-expanding clones (**Figure 5**, **Supplementary Table 4**). The effect is observed in genes involved in the *JAK/STAT* pathway, with *JAK1* (*P*=4.5x10-^7^), *JAK2* (*P*=0.0072), *STA1* (*P*=1.5x10^-7^), and *STAT2* (*P*=0.031) all downregulated amongst the expanding clones relative to non-expanding ones. In support of this, we see consistent downregulation of regulators of antiviral immune response *MX1* (*P*=5.2x10^-12^)*, ISG15* (*P*<2.22x10^-16^)*, and IFIT3* (*P*=2.5x10^-10^) amongst expanded cancer clones. Furthermore, the immunoproteasome genes *PSMB8* (*P*<2.22x10^-16^) and *PSMB9* (*P*<2.22x10^-16^) are downregulated, along with the associated regulatory protein *TAP1*. These results suggest that through mechanisms to dampen the IFN inflammatory response and reduce antigen presentation, expanding clones can avoid the host immune response, metastasise and expand.

### Up-regulation of cap-independent translational pathways is related to metastatic clonal behaviour

MRNA production in healthy cells involves cap-dependent translation mediated by the protein complex EIF4F. However, in cancer cells, cap-dependent translation is suppressed in favour of cap-independent translation, mediated by EIF4G1. This transition enables the sustained gene expression that promotes cell survival and profliferation^38^. Compared with healthy cells, we observe a higher ratio of *EIF4G1:EIF4F* as expected. However, we hypothesise that cap-independent translation pathway variation would impact metastatic cell phenotypes. To evaluate this, we compared the expression levels of cap-dependent translation genes between the expanded metastatic clones and non-expanding clones. The results display a clear pattern of upregulation of *EIF4G1* (*P*<2.22x10^-16^), coupled with the downregulation of genes *RPS2* (*P*<2.22x10^-16^), *RPS15* (*P*=1.1x10^-8^), *RPL23A* (*P*<2.22x10^-16^), and *RPL35A* (*P*<2.22x10^-16^) controlled by cap-dependent translation (**Figure 5**). Additionally, we observed the upregulation of genes involved in cancer cell proliferation and angiogenesis, with *HIF1A* (*P*=0.00012), *VEGFA* (*P*<2.22x10^-16^), and *PCBP1* (*P*=4.4x10^-7^) all upregulated in expanded versus non-expanded clones. Interestingly, whereas in other cancers, acquisition of somatic mutations has been linked to dysregulation of initiating factor genes^39^, we do not observe any evidence for that here (*EIF4A1, EIF4E3, EIF4G1, EIF4G2, EIF4EBP1*; **Figure 3**).

Our results identify the relationship between genomic phenotypes and clonal expansion into lymph node metastases, with expanding clones displaying a combination of dysregulated pathways favouring their survival. These can be summarised as the downregulation of cap-dependent translation and apoptosis, alongside the upregulation of cap-independent translation, co-expression with genes associated with cancer cell proliferation and metastatic processes, and evasion of immune response through the proteasome and IFN pathways.

### Evasion of immune cells within the primary tumour is a distinct cellular phenotype of metastatic clones

Our analysis of expanding clones that result in clonal dominance in the lymph node metastasis identified immune evasion through virally induced interferon and immunoproteasome pathways. We generated in situ spatial RNA-seq data from matched primary and lymph node biopsies to provide supportive evidence for this observation. We interrogated the relationship between cancer clonal subtypes and immune cell interactions within the cancer microenvironmental context. Numerous studies have highlighted the importance of the immune microenvironment within the primary site in influencing the metastatic potential of cancer cells^37,40^, and we aimed to test this in our cohort.

After classifying cancer clones and immune cell subtypes (**Methods**), we tested for the correlation in co-occurrence of pairwise combination of cells. We identified distinct differences between the T-cell correlation with expanding and non-expanding cancer clones (**Figure 6**). Expanding, metastatic clones predominately exhibit negative or neutral correlations (- 0.361<*R*<0.01) with T-cell subsets compared to more positive correlations in the non-expanding clones (-0.334<*R*<0.678). This observation is more pronounced within primary tumours. For example, the mean correlation in the primary between CD8 Exhausted cells (CD8Ex) and expanding and non-expanding clones (non-expanding = 0.599; expanding = - 0.12) is more disparate than in the lymph node (non-expanding = 0.295, expanding = 0.01). To evaluate this further, we tested for a significant difference in the median correlation between the expanding and non-expanding cancer clones across all patients. Adjusting for multiple testing, we show a significant difference in the co-occurrence of three T-cell subtypes between expanding and non-expanding cancer clones (**Figure 7A**). In the primary site, CD4 T-effector memory cells (CD4TEM) (*P*=0.0077), CD8 T-exhausted cells (CD8Ex) (*P*=0.0191), and innate lymphoid cells (ILC) (*P*=0.04) show a significant decrease in their correlation with expanding verses non-expanding cancer clones. CD4TEM cells also demonstrate a significant difference in correlation (*P*=0.021) with non-expanding clones compared to expanding clones in the lymph node site (**Figure 7A**). We observe a similar pattern of relationships in B-cell subsets (**Supplementary Figure 8**). The correlation analysis identified a distinct trend where expanding clones exhibit negative or neutral correlations with B-cells, in contrast to the behaviour observed in non-expanding clones, which have consistently strong co-occurrences. Summary correlations using mean correlation scores (**Figure 7B**) confirm this pattern. Notably, within the primary tumour, a substantial disparity exists in the correlations between expanding and non-expanding clones with B-memory cells and plasmablasts. Similarly, significant differences emerge in the correlations with B-memory (*P*=0.00048) and B-naïve (*P*=0.0065) cells within the lymph node. In the monocyte populations CD14 monocytes (CD14Mono), classical dendritic cells (cDC), plasmacytoid dendritic cells (pDC), a similar relationship to that observed in T and B-cell subsets (**Supplementary Figure 9**). The mean correlation scores (**Figure 7C**) identify a tendency to have positive correlations with monocytes and non-expanding clones, in contrast to the negative correlations noted with expanding clones. This trend is particularly pronounced in the lymph nodes, where significant differences between expanding and non-expanding clones exist within the CD14 monocyte (*P*=0.0091) and cDC subsets (*P*=0.0052).

**Figure 6.**
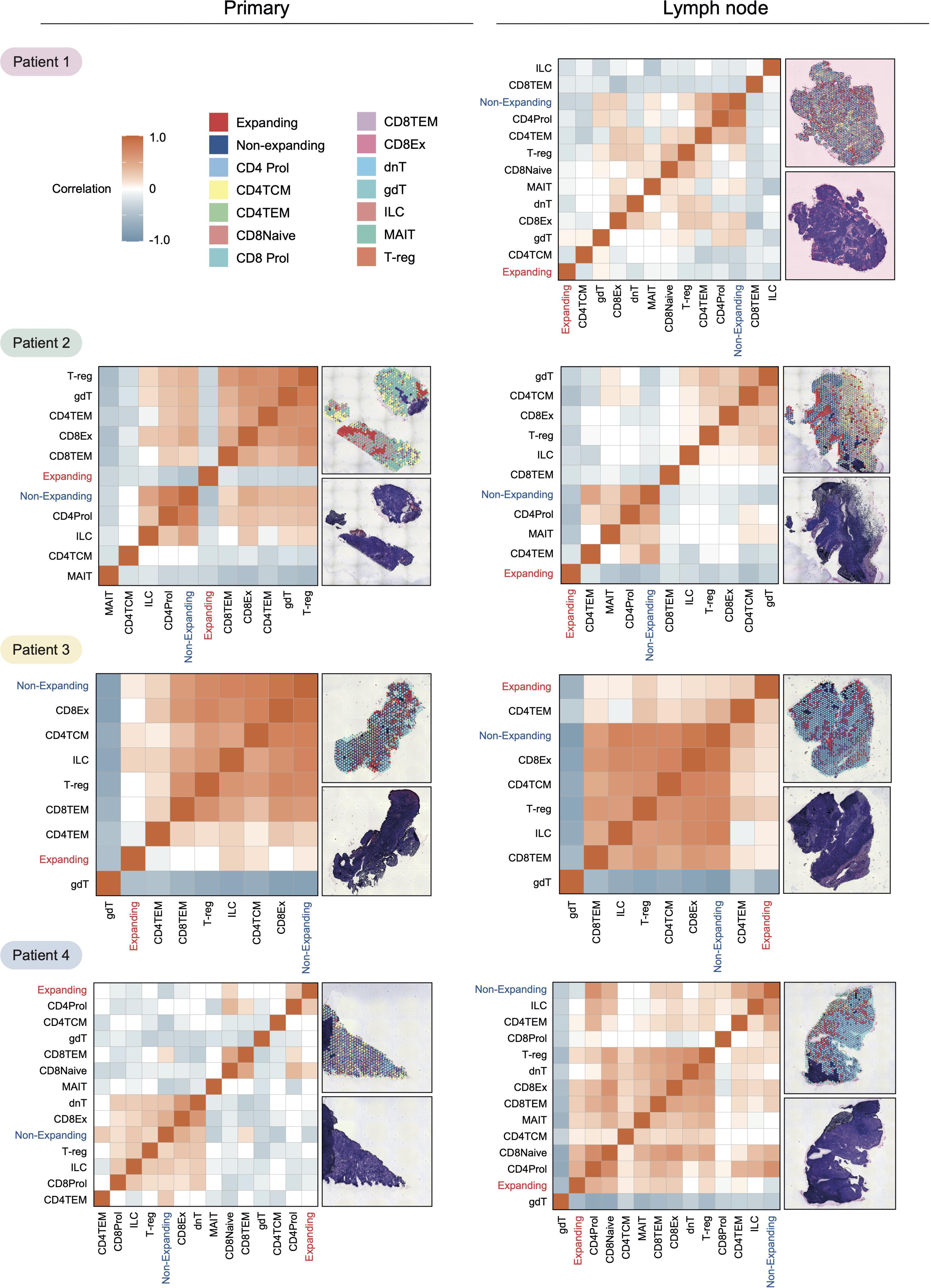
Relationship between expanding and non-expanding cancer clones and t-cell sub types. Spatial data was deconvoluted into cell type composition, and the colouration represents the estimated percentage of a cell’s contribution to a probe spot. Correlative analysis and pie chart plots are shown for the primary (left) and LN (right). CD8TEM = CD8 T effector memory cells, CD4Prol = CD4 proliferating cells, T-reg = T regulatory cells, MAIT = Mucosal-associated invariant T cells, dnT = Double negative t cells, CD8Ex = CD8 exhausted cells, gdT = gammadelta t cells, ILC = innate lymphoid cells.

**Figure 7.**
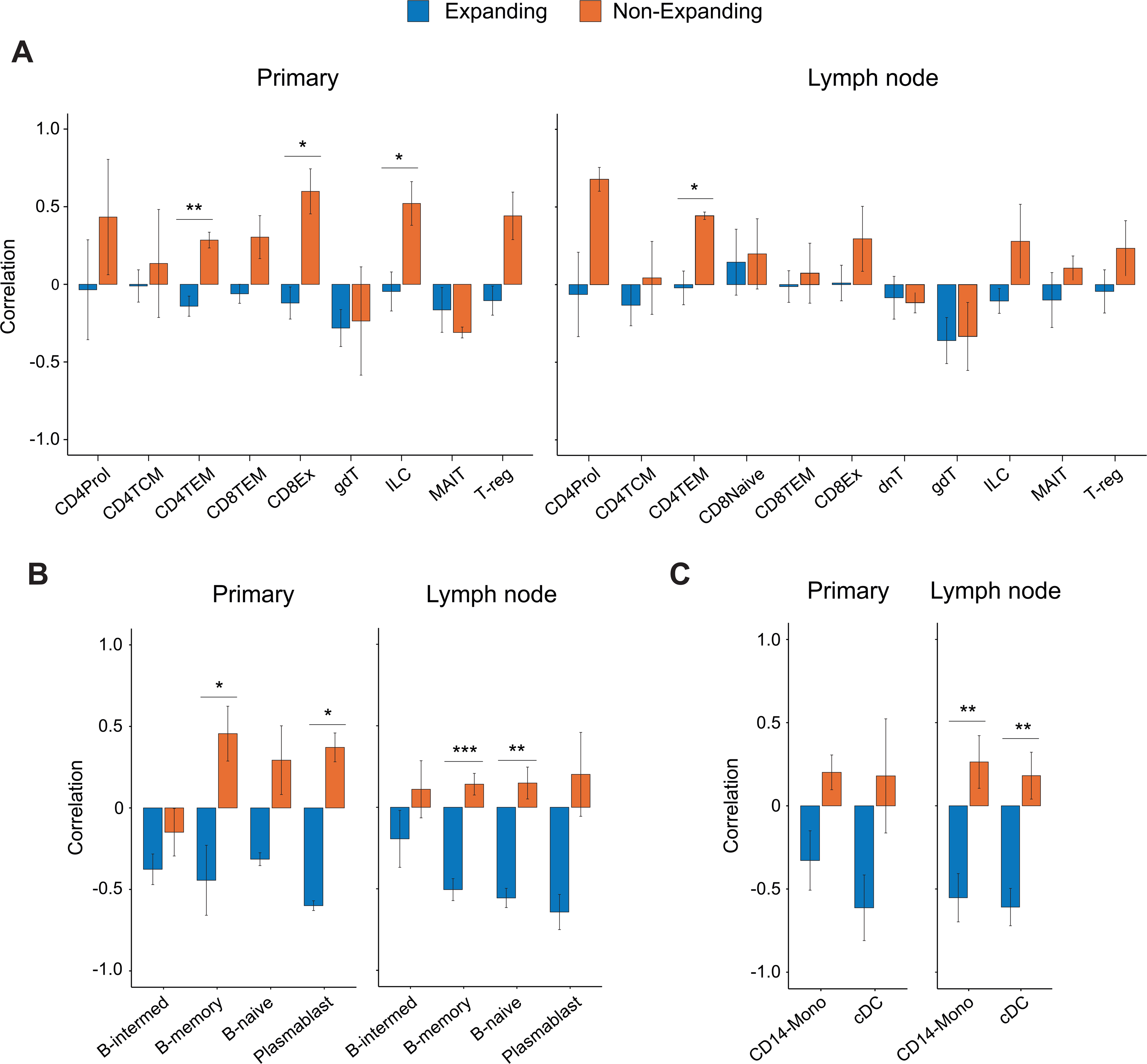
Relationship between immune cell and cancer clone co-occurrence across primary and lymph node sites. Co-occurrence correlations between expanding and non-expanding clones and immune cell subtype determined from spatial RNA in situ analysis. Significant differences were calculated and compared using t-tests, and study-wide significant relationships are displayed with a *.

The *in situ* data results demonstrate clone-specific immune evasion within the tumour microenvironment in metastasising cells. The observed dysregulation in the interferon and immunoproteasome pathways identified in differential gene expression analysis offers a credible explanation for this observed phenotype.

## DISCUSSION

It is widely acknowledged that cancer cells exhibit a range of diverse phenotypic properties^41^. Moreover, distinct cancer clones within tumours significantly influence the responses to anti-cancer treatments, subsequently impacting overall survival outcomes^42^. In locally advanced cancers, surgical resection aims to remove all the macroscopically identified cancer, with the addition of adjuvant therapies like radiation and chemotherapy to reduce recurrence rates by eradicating microscopic residual disease^43^. It has been shown that clones present in recurrent cancer are often sparsely represented in the primary tumour and that clonal complexity decreases in metastases^12,39^. While the mechanisms are not always resolved, it is clear that clonal selection occurs for cell phenotypes that enhance survival and metastatic processes. In the case of HPV+ oropharyngeal cancer, persistent HPV infection leads to alterations in the host immune response^30^, which likely results in significant selection pressure on cells, which shapes the tumour microenvironment^44^. This is evident when observing immune cell microenvironment changes through acute and chronic viral infection phases and pre-cancer and invasive cancer development with HPV infection^36^, highlighting the importance of examining cancer clones in their spatial context. Considering that HPV infection serves as the primary driver of carcinogenesis, our findings delineating metastatic clones are understandably distinct from those observed in the HPV-population^16^. Our initial results confirm observations presented in other cancers^45^, namely that there is reduced clonal complexity in metastases and that metastasising clones are sparsely represented in the primary tumour, confirming that clonal selection occurs. Through genomic analysis of cancer clones, we identified that the dysregulation of protein translation phenotypically defines expanding metastatic clones.

Given that protein translation dysregulation is seen in both acute^32^ and chronic^29,31^ viral infections and cancer^38^, we took a deeper look into the regulation of genes. We identified several important mechanisms that define expanding cancer clones in HPV+ oropharyngeal cancer. Our results suggest that virally induced translational relief is a mechanism that defines metastasising cancer clones in HPV+ oropharyngeal cancer. This observation aligns with prior work demonstrating HPV E6 protein is linked to translational control in infected fibrosarcoma cancer cell lines^29^, with E6 integration inducing the production of *EIF2AK2,* which in turn leads to phosphorylation of EIF2S1 and a decrease in protein translation and E6 production rapidly decreases. However, despite significant downregulation, the small amount of remaining E6 can initiate translational relief or rescue mechanisms. E6 can promote dephosphorylation of EIF2AK2 through stress kinase PPP1R15A, allowing protein translation to normalise and apoptosis to be inhibited^29^.

Furthermore, *PPP1R15A* is associated with tight regulation of the IFN-mediated anti-viral response^46^. In an acute viral infection, PPP1R15A is required to relieve protein translation inhibition to allow *IFN* and stress granule production in a pulsatile fashion. This allows the cell to restrict protein translation while releasing inflammatory mediators in periodic bursts. The dynamics of this process are different in chronic infection. When cells are exposed to stress, such as chronic viral infection^47^, PPP1R15A levels are continually elevated with fewer bursts, reducing stress granule release over time and creating an environment conducive to viral maintenance. Our results show that metastasising clones have increased expression of PPP1R15A and decreased levels of IFN-inducible genes, suggesting that these compensatory mechanisms favour viral replication and are critical in defining the metastatic phenotype. In addition to its role in viral response, *PPP1R15A* has previously been implicated in carcinogenesis and cancer growth. In mouse models of colorectal cancer, PPP1R15A potentiates carcinogenesis by enhancing *IL-6* production and STAT4 activation^48^. It has also been shown to promote autophagy in liver cancer^49^ and upregulate the RAC1-GTPase pathway in breast cancer^50^. This highlights a potential treatment target for the prevention of carcinogenesis in patients with chronic viral infection and established cancer.

Further perturbations of protein translation were identified in the metastatic cell phenotype, with the upregulation of cap-independent protein translation. Cap-independent mechanisms do not rely on the normal ribosomal machinery and, therefore, are not as susceptible to changes in functional subunit availability^51^. Thus, cancer clones that can utilise these cap-independent pathways to maintain protein translation may have an evolutionary advantage. Our data suggests that cap-independent translation maintains cancer-related proteins in the expanding metastatic clones. This observation is supported by cell-specific co-expression with cancer survival genes *VEGFA* and *HIF1A*. These results align with previous research in breast cancer models that have demonstrated how cap-independent mechanisms selectively translate mRNAs that encode proteins critical for cancer cell survival like VEGFC, HIF1A and BCL2^52,53^. Previous work shows that lung cancers rely on cap-independent mechanisms to maintain cell replication and that EIF4G1 is an adverse prognostic feature^38^. Given that cap-independent translation appears to be a mechanism underlying the selection of HPV+ oropharyngeal cancer metastatic clones, we propose this may be a useful biomarker to identify patients with poor prognostic disease.

Immune cell evasion was identified as a defining feature of metastatic cancer clones both as a cellular phenotype and through analysis of the tumour microenvironment. For HPV infection to persist, viral proteins E5, E6 and E7 mediate anti-inflammatory and immune-evading functions. For example, in human keratinocytes, E5 can actively suppress *IFN* and IFN-stimulated gene expression, key components of the innate anti-viral response^54^. Here, we show that the downregulation of immunoproteasomes is an important component of immune evasion in the expansion and metastasis of clones in HPV+ oropharyngeal cancer. The immunoproteasome is involved in antigen presentation and is a critical step in the inflammatory response to viral infection and cancer. E5 HPV protein is a powerful negative regulator of the anti-viral immune response by limiting the MHC I antigen repertoire via suppression of the immunoproteasome^30^, allowing infected cells to escape host immune surveillance^55,56^. The importance of the immunoproteasome in antigen recognition is further demonstrated, with high expression being a positive predictive biomarker of immune checkpoint inhibitor blockade response^57^. Our data shows that immune evasion is a second phenotypic feature of metastatic clones. Although HPV+ oropharyngeal cancer is considered an immunotherapy-sensitive cancer, the response is not universal. In the metastatic setting, survival is less than two years despite checkpoint inhibitor use. Ineffective antigen presentation via the immunoproteasome may be a desirable therapeutic target in HPV+ oropharyngeal cancer, given that HPV infection and carcinogenesis are dysregulators of this process. Restoration of the immunoproteasome by IFN-γ has been shown to improve immune system recognition of renal cancer cells^58^ and cervical cancer^59^. Interestingly, when this approach was used in head and neck cancer^60^, highly heterogeneous responses were seen *ex vivo*. This heterogeneity could be explained by the mixture of HPV+ and HPV-cancers analysed.

Challenges for managing HPV+ oropharyngeal cancer include the lack of reliable prognostic markers and paucity of druggable targets. This limits the ability to personalise treatment decisions in curable disease and offer effective treatment options in the metastatic setting. We have identified mechanisms defining the metastatic clonal phenotype unique to HPV+ oropharyngeal cancer. Given the aetiology of HPV-related cancer, it is expected that many of the cellular mechanisms are shared with those observed in chronic viral infection. *EIF4G1* expression may be helpful as a prognostic marker, which could be used to select patients more likely to metastasise and benefit from more aggressive curative therapies. Finally, the immunoproteasome likely plays a significant role in immune evasion in metastasising clones. Further investigation of immunoproteasome proteins as predictive biomarkers for checkpoint inhibitor response and direct targeting as a therapeutic strategy should be undertaken.

## Acknowledgements

We acknowledge the patients and their families.

This research was funded by Cure Cancer Australia, St Vincent’s Clinic Research Foundation and The Garvan Institute of Medical Research Foundation.

## Author Contributions

V.C. designed the biobanking study and research project, conducted the experiments, analysed data, and wrote the manuscript. W.M. optimised *Numbat* code and assisted with data analysis and graphics. R.M. conducted biobanking, ethics management and conducted experiments. J.A. conducted immune cell subtyping. D.N. and H.A. conducted de-multiplexing and pre-processing of single cell data. A.S. assisted with data analysis, storage and pre-processing. P.K. conducted biobanking. E.S. conducted biobanking and experiments. A.M. conducted biobanking and experiments. D.K. conducted experiments. P.F, B.L and J.C. consented patients and retrieved specimens. P.E. specimen preparation, cut up and histopathological analysis. R.G. managed the clinical aspects, consented patients, retrieval of specimens. J.P. designed experiments, assisted with data analysis, and wrote the manuscript.

## Declaration of Interests

Nil

## Methods

### RESOURCE AVAILABILITY

#### Lead contact

Further information and requests for resources and reagents should be directed to and will be fulfilled by the lead contact, A/Prof Venessa Chin

## Materials availability

This study did not generate any new unique reagents

## Data and code availability

All single-cell RNA-sequencing data generated by this study have been deposited in the _______ (https://). The data can be accessed under the accession number______. All other data supporting the findings of this study and Code used for all processing and analysis are available from the corresponding author upon reasonable request.

## EXPERIMENTAL MODEL AND STUDY PARTICIPANTS DETAILS

### Ethics statement

This project received ethics approval from the St Vincent’s Hospital Human Research Ethics Committee (HREC/16/SVH/329) in accordance with the ethical standards laid down in the Declaration of Helinski. Written informed consent was obtained per institutional guidelines.

### Patient cohort

Patients were an average age of 55 and compromised 75% men (n=3) and 25% women (n=1). Newly diagnosed, previously untreated patients with HPV+ Oropharyngeal squamous cell carcinoma were selected if they presented with lymph node metastases at St Vincent’s Hospital Sydney, and proceeded to surgery for their tumour.

## METHOD DETAILS

### Tissue sample processing

Within 30 minutes of surgical resection, tumour samples were tumour banked. For Visium, half of the tumor sample was cut off and snap frozen in O.C.T (Tissue-Tek, CAT# 4583) on dry ice. After freezing, samples were moved to -80°C for long-term storage. The remaining sample was cut into 2mm x 2mm chunks with a scalpel blade. For single cell analysis, 2-3 tumour chunks were placed into a cryovial with 90% Fetal Calf Serum (FCS) + 10% DMSO. Cryovials were placed into a CoolCell in a ^-^80°C freezer overnight, then transferred to Vapour Phase for long-term storage.

### Blood processing

Blood was collected in 10mL BD Vacutainer EDTA tubes (Becton Dickinson, CAT# 367525). The sample was processed for plasma and buffy coat as previously described ^61^. Samples were stored at -80 °C.

### SNP Genotyping

DNA was extracted from Buffy Coat samples using the QIAamp DNA Blood Mini Kit (Qiagen, CAT# 51106). Genotyping was performed using the UK Biobank Axiom array (ThermoFisher Scientific, CAT# 902502) at the Ramaciotti Centre for Genomics. Imputation was performed using the Michigan Imputation Server, with Minimac4 and the Haplotype Reference Consortium (HRC) panel.

### Preparation of Single Cell Suspensions

The presence of cancer tissue was confirmed on haematoxylin and eosin staining before sequencing. Tissue samples were defrosted in a 37°C water bath. Contents were transferred to a 6 cm culture dish and washed with RMPI-1640 (Gibco, CAT# 11875093) + 10% FCS. Samples were moved to a 2mL Lo-Bind Eppendorf containing 100uL of RPMI-1640 and mechanically dissociated using scissors and snips. Samples were chemically dissociated with the Miltenyi Tumour Dissociation Kit (Human) (Miltenyi Biotech, CAT# 130-095-929) according to the manufacturer instructions. Samples were incubated for 20-30 minutes on a shaking incubator at 37°C until all tumour chunks were dissociated. Sample solution was passed through a 100um cell strainer into a FACS tube with RMPI-1640 + 10% FCS. Cells were counted and viability assessed using Trypan Blue. If samples had <80% cell viability then dead cells were removed with the EasySep Dead Cell Removal (Annexin V) Kit (StemCell Technologies, CAT# 17899) according to manufacturers instructions. Samples were pooled equally (primaries pooled together and lymph nodes pooled together) to a final concentration of 1500 cells per uL in PBS + 10% FCS. Samples were passed through a 70uM cell strainer.

### Single-Cell Capture and cDNA Library Preparation

The 10X Genomics Chromium instrument (10X Genomics) was used to partition viable cells with barcoded beads and cDNA from each cell was prepared using the 10X Genomics Single Cell 3’ Library, Gel Bead and Multiplex Kit (v3) (CAT# 10X-1000075) as per the manufacturer’s instructions. Patient samples were pooled and multiplexed. Cell numbers were optimised to capture approximately 5000 cells per individual sample, and 20000 cells in the total pool. Samples were loaded onto a 10X Genomics Chromium Single Cell Chip Kit B (CAT# 10X-1000153). Libraries were generated with the 10X Genomics Chromium Single Cell 3’ Library construction kit (v3) (CAT# 10X-1000078)

### scRNA-seq Data Processing

#### Library sequencing

The libraries were sequenced at the Ramaciotti Centre for Genomics on an Illumina NovaSeq 6000 (NovaSeq Control Software v 1.7.0 / Real-Time Analysis v3.4.4) using a NovaSeq S4 200 cycles kit (Illumina, 20482067 as follows: 28bp (Read 1), 91bp (Read 2) and 8bp (Index). A median sequencing depth of 30,000 reads/cell was targeted for each sample. The sequencer generated raw data files in binary base call (BCL) format. The BCL files were demultiplexed and converted to the FASTQ file formats using Illumina Conversion Software (bcl2fastq v2.19.0.316).

### Bioinformatics pipeline

The cellranger -v (3.1.0) count pipeline was used for alignment, filtering, barcode, and UMI counting from FASTQ files. The pipeline was executed on a high-performance cluster with a 3.10.0-1127.el7.x86_64 operating system. RNA sequencing reads were mapped to the human GRCh38 transcriptome and HPV 16 and 18 transcriptome ^62,63^ using the STAR aligner ^64^.

**DropletQC –** To distinguish between true cells and cell-free ambient RNA, DropletQC. Empty drops were excluded from the analysis^65^.

**Demultiplexing** – SNP data was generated for each individual as described above. Multiplexed patient pools were demultiplexed using Demuxafy ^66^. Doublets were identified and removed.

**Single Cell Data –** The aggregated single-cell gene expression data generated by CellRanger was used as the input for the Seurat analysis software 4.0 ^67^. Expression levels for each transcript were determined using the number of unique molecular identifiers assigned to the transcript. Quality control and filtering steps were performed to remove outlier genes and cells. Cells with an expression greater than 25% of mitochrondrial genes were removed. Cell-cell normalisation was performed using a regularised negative binomial regression (sctransform)^68^. Principal component analysis was performed on the filtered and normalised gene expression matrix, and the first 15 principal components that explained the majority of the variance in the data were retained. Uniform manifold approximation and projection for dimension reduction (UMAP)^OBJ^ plots were generated to visualise the gene expression patterns in each cell.

**Integration of data –** Seurat was used to identify common anchors between datasets to integrate primary and lymph node data for each individual into the same object ^70^.

### CNV Estimation and identification of malignant cells

Clonal architecture was inferred using the Numbat package^25^. Cancer cells from each individual were examined separately, with cells from both the primary and lymph node metastasis grouped, using the individual’s immune cells as a normal reference. The default running parameters were used for all samples except patient 1. In this sample, the clones did not become stable until the LLR reached 400. This was thought to be due to the high number of malignant cells in this sample. Cancer cells with similar CNV profiles were grouped in a clonal group.

**Immune cell classification –** Cancer cells were identified using *EPCAM* expression. The immune cells were subtyped using automated methods using a reference dataset for immune cells^71^. This reference dataset does not include exhausted CD8+ t-cells or macrophages, so these were added manually (Figure 1C). CD8+ T-central memory cells (CD8 TCM) were identified using the reference dataset. Markers of cytotoxicity (*GNLY*, *GZMK*, *TNF*, *FASLG*, *PRF1*, *IL2*, *IFNG*) and exhaustion (*LAG3*, *BTLA*, *CTLA4*, *PDCD1*, *HAVCR2*) were used to classify the CD8 TCMs further. One sub-population of CD8 TCMs expressed high cytotoxicity and exhaustion markers and were labeled as CD8TCM-exhausted cells. For macrophages, the monocyte population was identified using the reference dataset. Expression counts within the monocytes of *FCGR2A* were calculated. Any cells expressing FCGR2A >2x the mean expression was assigned as a macrophage. After cell filtering, normalisation, and cell classification we had 14157 cancer cells 36324 immune cells and 308 stromal/endothelial cells for analysis.

**Differential Gene Expression Analysis –** To examine the phenotype of the expanding clones, the cancer clones were grouped into transcriptionally similar clusters with the assigned clones overlayed. The non-expanding clones in the primary were compared with the expanding clones in the lymph node for differential gene expression analysis. Differential gene expression analysis was performed with the “FindAllMarkers” function in Seurat^67^. Calculations were corrected for the patient pool. Where there were transcriptionally and clonally distinct clusters, multiple comparisons were made per patient. Patient 1 and patient 3 had two comparisons done each. Patient 1 had two transcriptionally distinct clusters in the primary - “P-cluster A”, made up of clone 4, and “P-cluster B”, made up of clone 6. These were each separately compared to the LN cluster, denoted 1A and 1B in Figure 4B. Similarly, in patient 3, the primary cluster was compared to each of the LN clusters, “LN-cluster A”, made up of primarily clone 3 and “LN-cluster B”, primarily made up of clones 4 and 5 (denoted comparison 3A and 3B in Figure 4B). DEG analysis was performed for each patient and analysed with ReactomePA^72^ for enriched pathways. Custom code was developed for plots in supplementary figures 2-6.

### Visium Spatial Gene Expression

Following the manufacturer’s instruction, O.C.T embedded tissue samples were processed using the Visium Spatial Gene Expression slide and reagent kit (10X Genomics, PN-1000186). Briefly, 10 μm cryosections were placed into the capture areas of the Visium slide. Tissue morphology was assessed with H&E staining and imaging using a Leica DM6000 Power

Mosaic microscope equipped with a 20x lens (Leica). The imaged sections were then permeabilized for 18 minutes using the supplied reagents. The permeabilization condition was previously optimised using the Visium Spatial Tissue Optimization reagent kit (10X Genomics, PN-1000192), and Visium Spatial Tissue Optimization slide kit (10X Genomics, PN-1000191). After permeabilization, cDNA libraries were prepared using the Library Construction Kit (10X Genomics, PN-1000190), checked for quality and sequenced on a NovaSeq 6000 platform (Illumina, US). Sequencing targeted 50,000 reads per occupied spot on the Visium slide. Around 300 million pair-ended reads were obtained for each tissue section. Read 1, i7 index and Read 2 were sequenced with 28, 8 and 91 cycles respectively.

### Visium spatial transcriptomics data processing

Three out of four patients had suitable tissue for spatial sequencing for both the primary and lymph node; however, in patient one, only the lymph node sample had sufficient tissue. Reads were demultiplexed and mapped to the reference genome GRCh38 using the Space Ranger Software v1.0.0 (10X Genomics). Count matrices were loaded into the Seurat v3.2 (https://github.com/satijalab/seurat/tree/spatial) and STutility (https://github.com/jbergenstrahle/STUtility) R packages for all subsequent data filtering, normalisation, filtering, dimensional reduction and visualisation. All spatial spots determined to be over tissue regions by Space Ranger were kept for subsequent analysis. Poor quality tissue locations were then filtered out based on a cut off of 500 unique genes. Genes detected in more than 10 locations were also kept for analysis. Data normalisation was performed on independent tissue sections using the variance stabilizing transformation method implemented in the SCTransform function in Seurat. We applied non-negative matrix factorization (NMF) to the normalised expression matrix using the STutility package (nfactors = 20). NMF reduction was then used for clustering using Seurat with all 20 factors as input (RunUMAP, FindNeighbors and FindClusters functions).

**Deconvolution of Visium Spots** – for each patient, the single cell dataset was used as a reference for deconvolution of the Visium spots. The cell subtype signatures from annotated single cell data is used to determine the cell composition of each Visium spot using a seeded NMF regression^74^. A weighted combination of cell types which reside in each spot are calculated and visualised as a pie chart. With these weighted combinations, correlation analyses were performed, examining the likelihood of a certain cell-types co-locating with one another.

### Correlation plots

Mean correlation scores between cancer clones and cell types of interest were calculated for all individual primaries and LN separately. Bar graphs were generated using ggbarplot^73^, and statistical significance was determined using a t-test.

## QUANTIFICATION AND STATISTICAL ANALYSIS

### Statistical analysis performed and software used

Figure 2 – statistical significance was determined using Fisher exact test in rstatix package^75^ Figure 5 – statistical significance was determined using a t-test using the stat_compare_means function in ggplot2.

P-values are reported with asterix: * <0.05, ** <0.01, ***<0.001

### Key Resources Table

| REAGENT or RESOURCE | SOURCE | IDENTIFIER |
| --- | --- | --- |
| Biological samples |  |  |
| Fresh surgical tumor resections | The St Vincent's Head and Neck Cancer Biobank | Ethics # HREC/16/SVH/329 |
| O.C.T embedded tumor tissue blocks | The St Vincent's Head and Neck Cancer Biobank | Ethics # HREC/16/SVH/329 |
| Buffy Coat, prepared from whole blood | The St Vincent's Head and Neck Cancer Biobank | Ethics # HREC/16/SVH/329 |
| Critical commercial assays |  |  |
| Chromium Next GEM Single Cell 3' Library, Gel Bead and Multiplex Kit v3 | 10X genomics | 10X-1000075 |
| Chromium Next GEM Single Cell Chip Kit B | 10X genomics | 10X-1000153 |
| Chromium Single Cell 3' Library construction kit v3 | 10X genomics | 10X-1000078 |
| Visium Spatial Gene Expression Slide and Reagent Kit | 10X genomics | PN-1000186 |
| Library Construction Kit | 10X genomics | PN-1000190 |
| Miltenyi tumor dissociation kit, human | Miltenyi | 130-095-929 |
| EasySep Dead Cell Removal (Annexin V) Kit | StemCell Technologies | 17899 |
| QIAamp DNA Blood Mini Kit | Qiagen | 51106 |
| Deposited data |  |  |
| Single cell RNA-seq data? | This paper |  |
| Visium data? | This paper |  |
| Software and algorithms |  |  |
| Cell Ranger v 3.1.0 | 10X Genomics | https://software.10xgenomics.com/ |
| STAR aligner | Dobin et al., 2013 | https://github.com/alexdobin/STAR |
| DropletQC | Muskovic and Powell, 2021 | https://powellgenomicslab.github.io/DropletQC/ |
| Demuxafy | Neavin et al., 2022 | https://github.com/drneavin/Demultiplexing_Doublet_Detecting_Docs |
| Seurat 4.2.2 | Satija et al., 2015 | https://github.com/satijalab/seurat |
| scPred | Alquicira-Hernandez., 2019 | https://github.com/powellgenomicslab/scPred |
| Numbat | Gao et al., 2022 | https://github.com/kharchenkolab/numbat |
| karyoploteR | Gel et al., 2017 | https://bernatgel.github.io/karyoploter_tutorial/ |
| ReactomePA | Yu et al., 2016 | https://github.com/YuLab-SMU/ReactomePA |
| ggplot2 | Wickham. 2016 | https://ggplot2.tidyverse.org/ |
| SPOTlight | Elosua-Bayes et al., 2021 | https://marcelosua.github.io/SPOTlight/ |
| rstatix | Kassambara et al.,<br>2023 | https://rpkgs.datanovia.com/rstatix/ |

## Supplemental information titles and legends

**Supplementary Table 1.** Patient characteristics and survival outcomes

**Supplementary Table 2.** Differential gene expression analysis for the comparisons in Figure 4. Each comparison is listed separately.

**Supplementary Table 3.** Gene enrichment analysis results for each comparison in Figure 4. Each comparison is listed separately.

**Supplementary Table 4.** Mean expression levels of genes shown in Figure 5

**Supplementary Figure 1.** Patient 1A. The top ten enriched pathways, revealing the direction and extent of gene expression changes

**Supplementary Figure 2.** Patient 1B. The top ten enriched pathways, revealing the direction and extent of gene expression changes

**Supplementary Figure 3.** Patient 2. The top ten enriched pathways, revealing the direction and extent of gene expression changes

**Supplementary Figure 4.** Patient 3A. The top ten enriched pathways, revealing the direction and extent of gene expression changes

**Supplementary Figure 5.** Patient 3B. The top ten enriched pathways, revealing the direction and extent of gene expression changes

**Supplementary Figure 6.** Patient 4. The top ten enriched pathways, revealing the direction and extent of gene expression changes

**Supplementary Figure 7.** Functional protein association network from patient comparison 1A

**Supplemental Figure 8.** S**p**atial **data and correlative analysis for cancer clones and b-cell subsets.** Correlative analysis and pie chart plots are shown for the primary (left) and LN (right). B-intermed = b intermediate cells.

**Supplemental Figure 9.** S**p**atial **data and correlative analysis for cancer clones and monocyte subsets.** Correlative and pie chart plots are shown for the primary (left) and the LN (right). CD14 Mono = CD14 positive monocytes, cDC = classical dendritic cells, pDC = plasmacytoid dendritic cells

## Notes

### Competing Interest Statement

The authors have declared no competing interest.

