## Supplementary figures and images for "Protein Translation Dysregulation and Immune Cell Evasion Define Metastatic Clones in HPV-related Cancer of the Oropharynx"

### Supp Fig 1

##### Supplementary Figure S1

Patient 1A

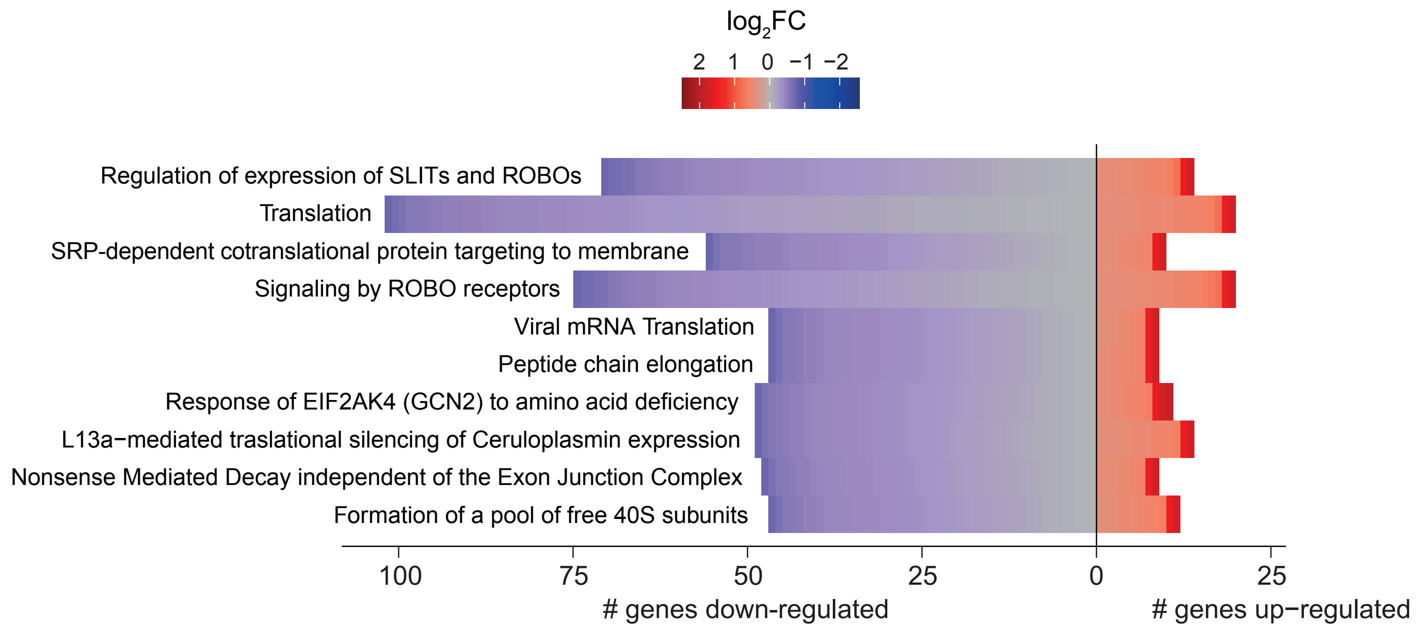

### Supp Fig 2

#### Supplementary Figure S2

Patient 1B

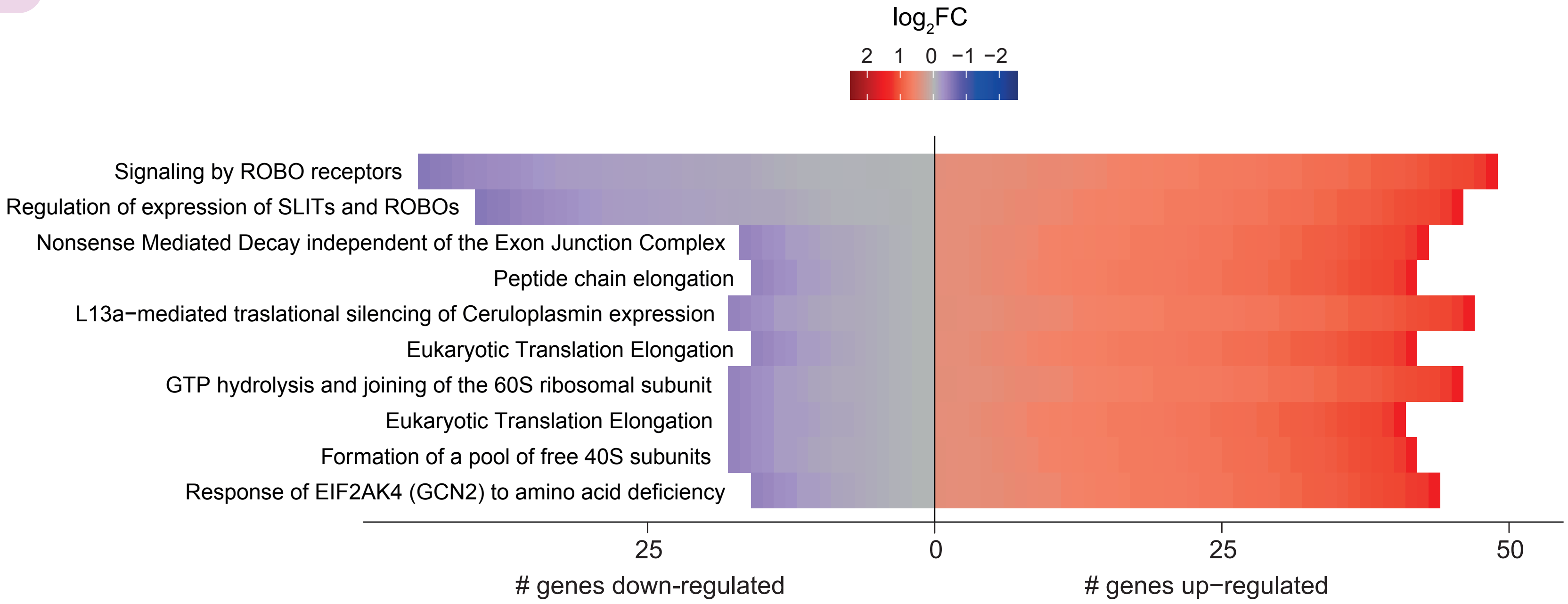

### Supp Fig 3

#### Supplementary Figure S3

Patient 2

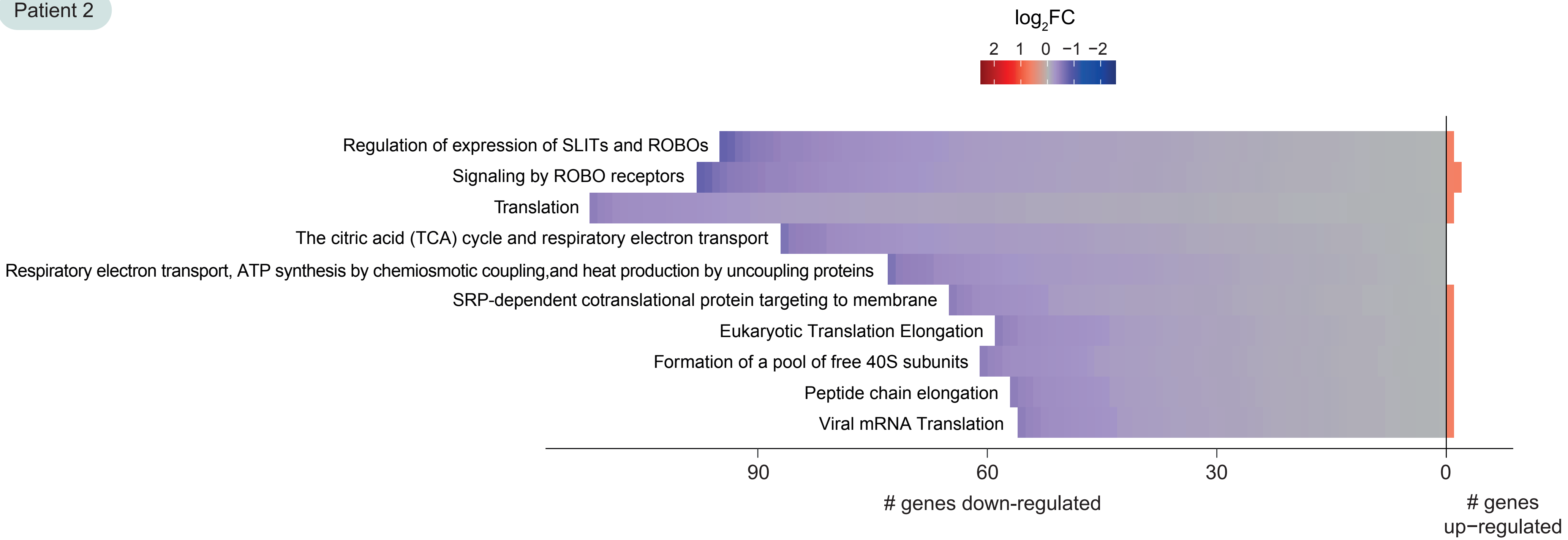

### Supp Fig 4

##### Supplementary Figure S4

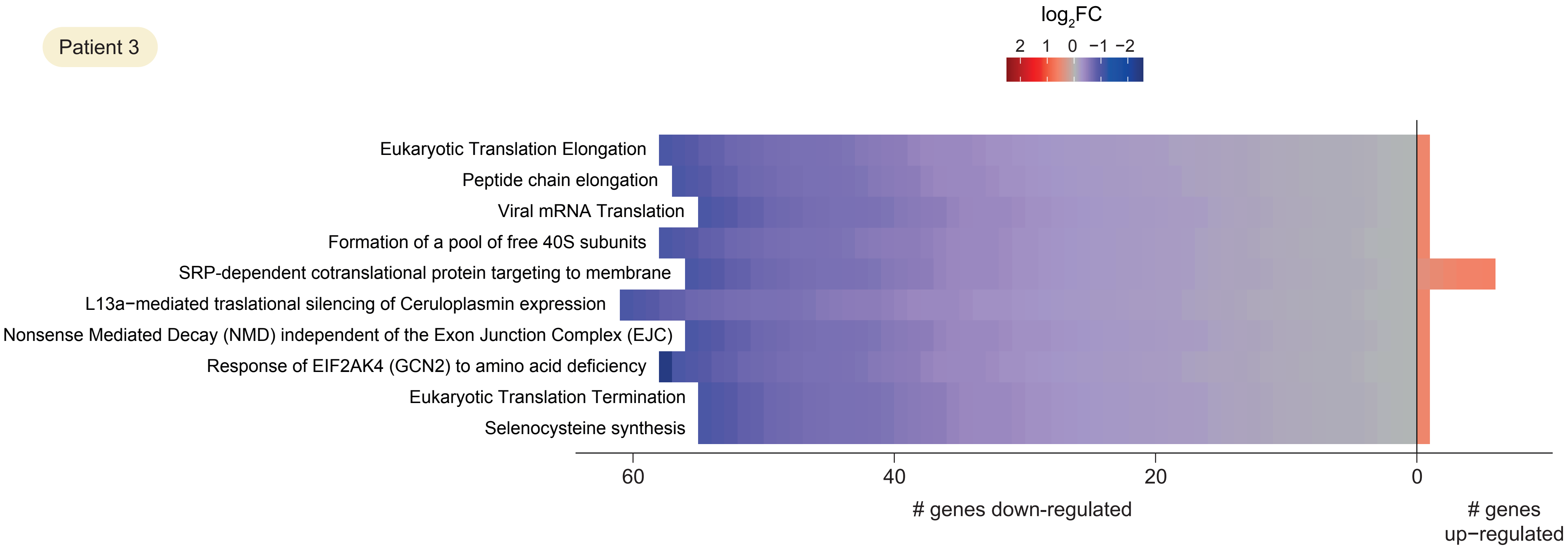

### Supp Fig 5

#### Supplementary Figure S5

Patient 3B

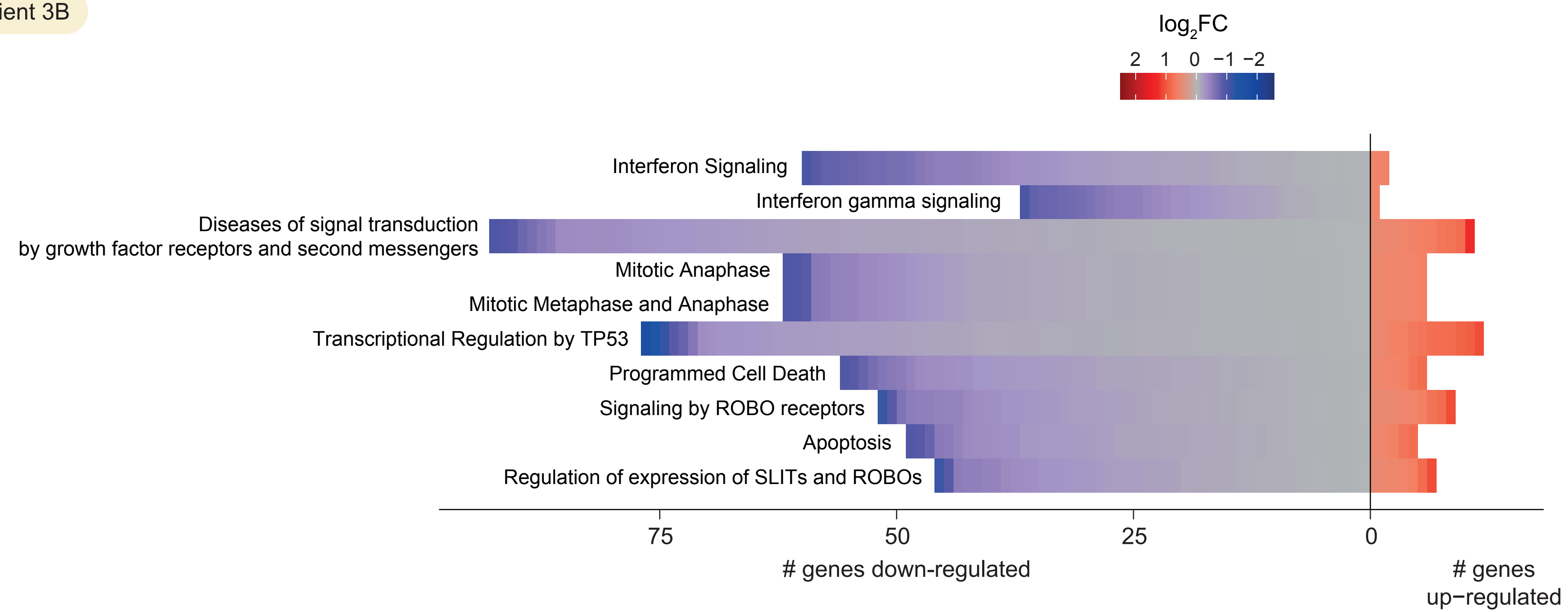

### Supp Fig 6

##### Supplementary Figure S6

Patient 4

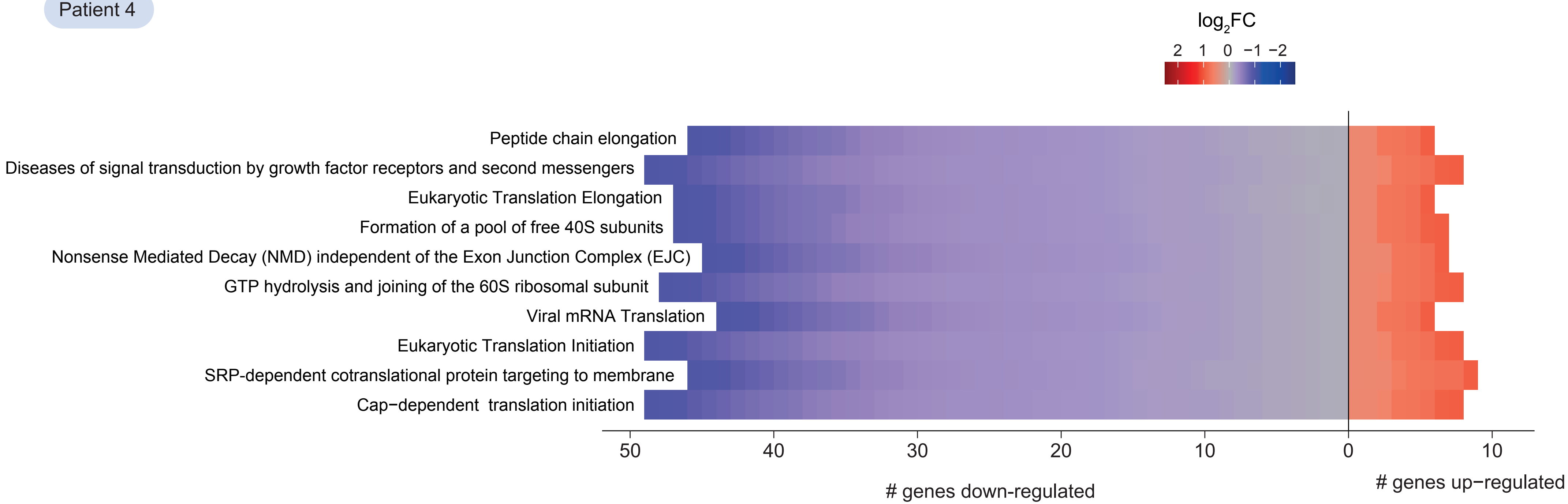

### Supp Fig 7

##### Supplementary Figure S7

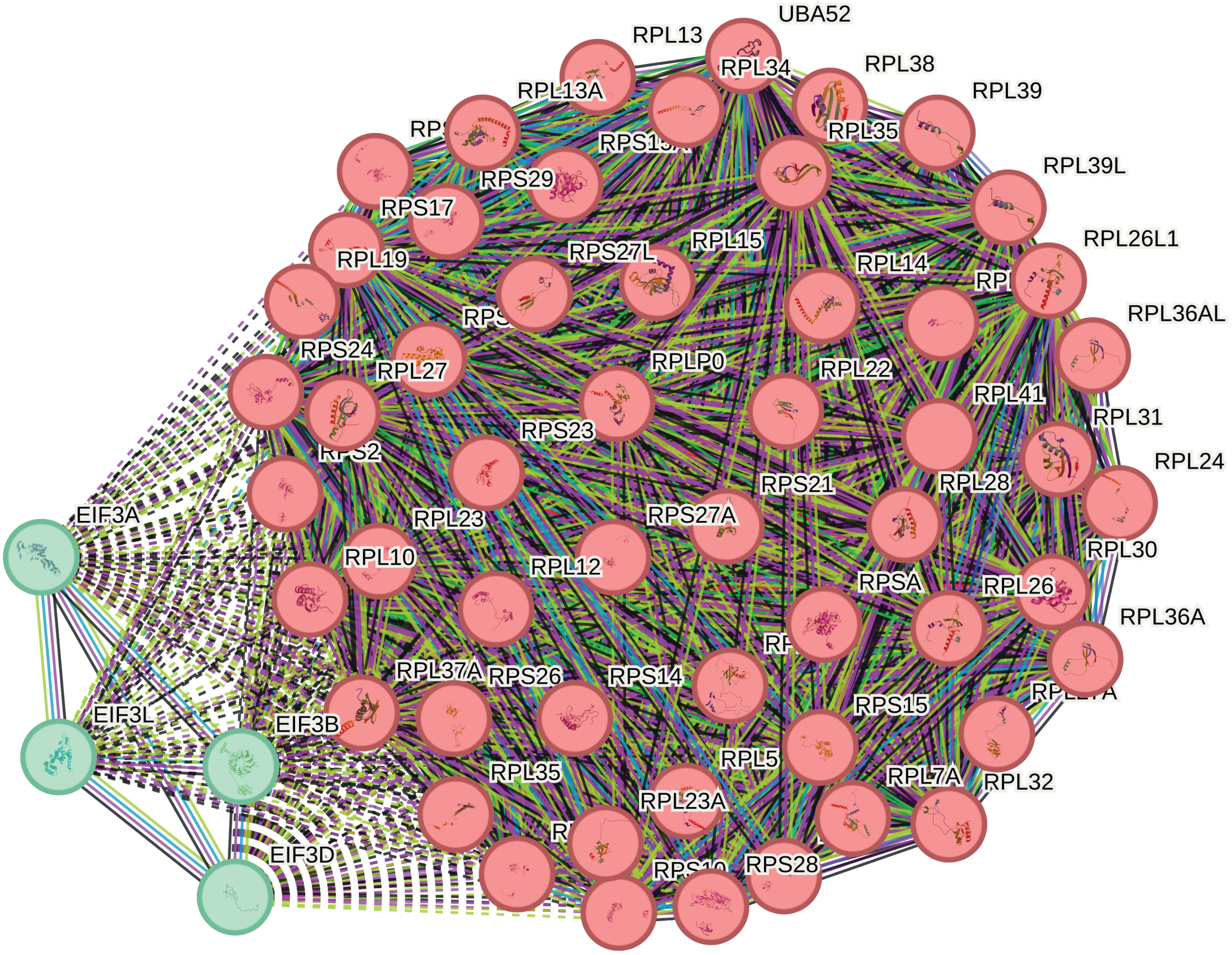

### Supp Fig 8

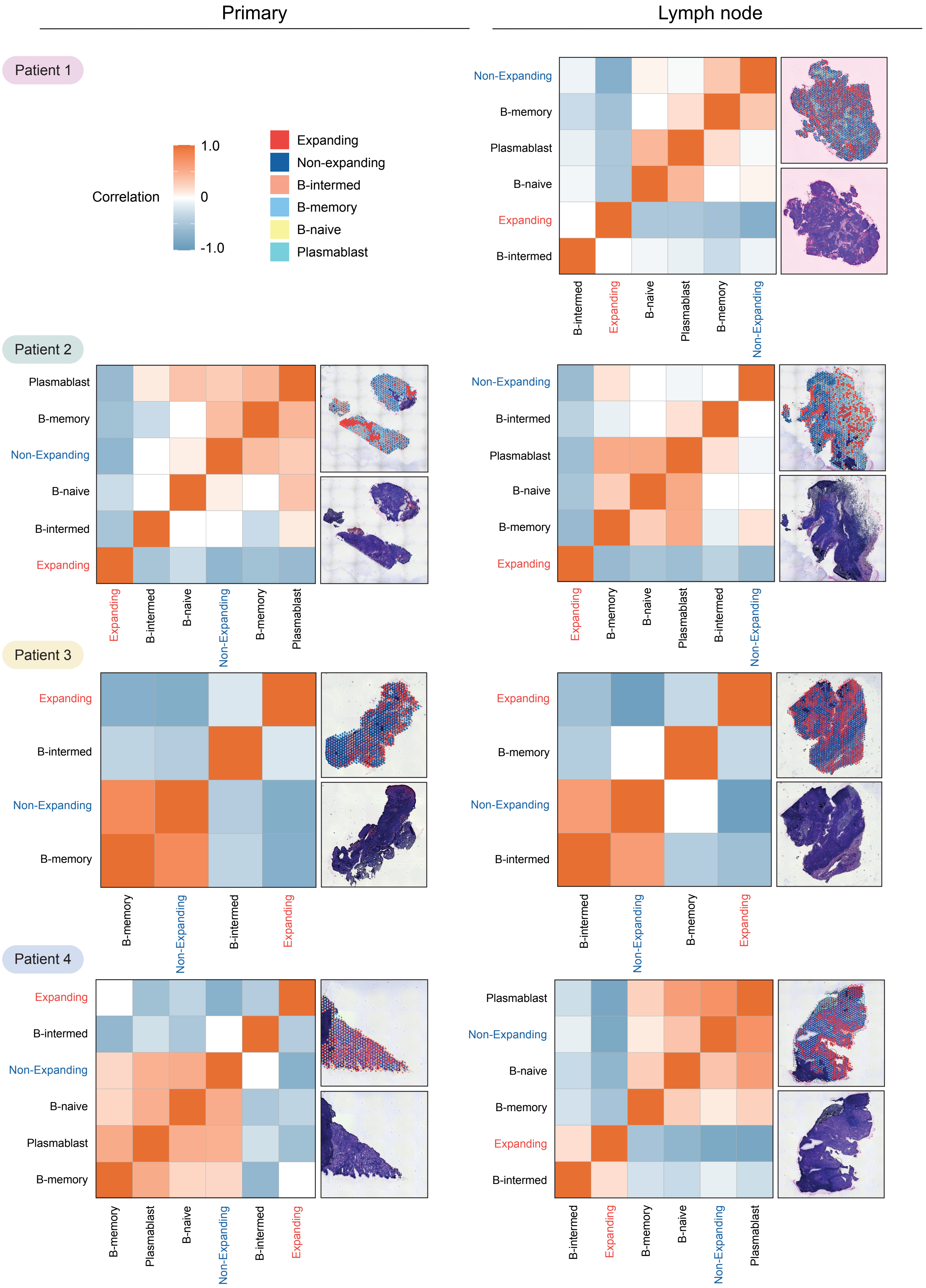

### Supp Fig 9

#### Supplementary Figure S8

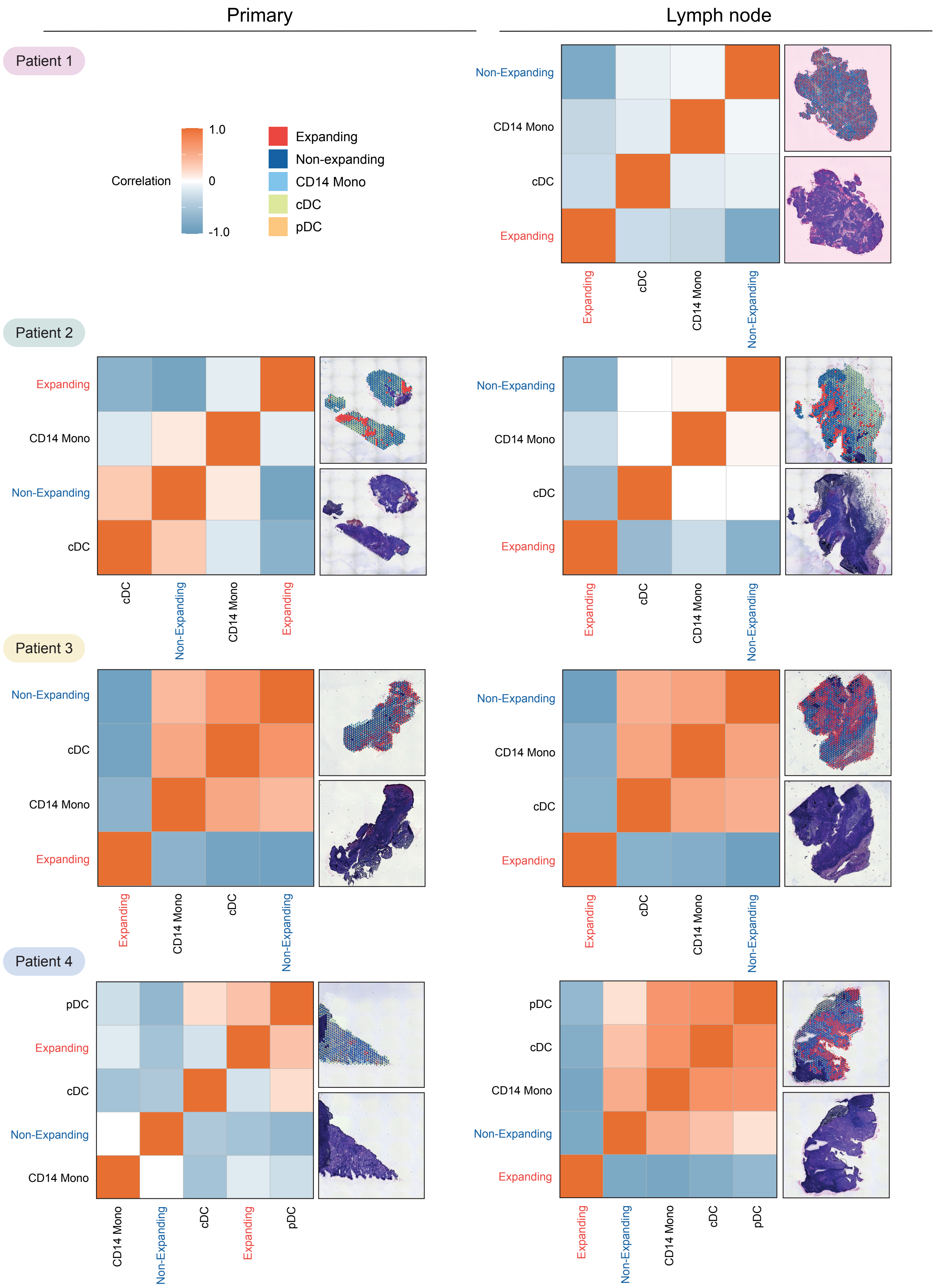
