## Supplementary material for "Protein Translation Dysregulation and Immune Cell Evasion Define Metastatic Clones in HPV-related Cancer of the Oropharynx": Supp Table 1

### Supplementary Table S1

| Patient characteristics | |
|---|---|
| **Median Age** | 61 |
| **Male (%)** | 86 |
| **Smoking status (%)** | |
| Former | 71 |
| Current | 29 |
| **Primary site (%)** | |
| Tonsil | 86 |
| Base of tongue | 14 |
| **T stage (%)** | |
| T1 | 43 |
| T2 | 57 |
| **N stage (%)** | |
| N1 | 86 |
| N2 | 14 |
| **Positive margins (%)** | 14 |
| **Extra-nodal extension (%)** | 29 |
| **Adjuvant radiotherapy (%)** | 57 |
| **Adjuvant chemoradiotherapy (%)** | 29 |
| **Median follow up (months)** | 40 |
| **Recurrence (%)** | 28 |
| **Death (%)** | 14 |
