## Supplementary material for "Protein Translation Dysregulation and Immune Cell Evasion Define Metastatic Clones in HPV-related Cancer of the Oropharynx": Supp Table 4

### Supplementary Table S2

| | Mean Expression | | |
|---|---|---|---|
| Gene | Non-Expanding | Expanding | P-value |
| EIF4A1 | -0.005 | 0.06 | 0.1013 |
| EIF4E | -0.155 | -0.0903 | $1.5 \times 10^{-13}$ |
| EIF4G1 | -0.156 | 0.07529 | $<2.22 \times 10^{-16}$ |
| PDCD4 | -0.1756 | -0.103 | $5.3 \times 10^{-14}$ |
| EIF4EBP1 | 0.076 | -0.0367 | $2.1 \times 10^{-6}$ |
| EIF2AK2 | 0.13406 | -0.0645 | $<2.22 \times 10^{-16}$ |
| EIF2S1 | 0.112 | -0.054 | $<2.22 \times 10^{-16}$ |
| PPP1R15A | -0.466 | -0.343 | $<2.22 \times 10^{-16}$ |
| PPP1CA | -0.0509 | 0.0245 | $7.9 \times 10^{-6}$ |
| PPP1CB | -0.148 | 0.071 | $<2.22 \times 10^{-16}$ |
| PPP1CC | 0.077 | -0.037 | $5 \times 10^{-10}$ |
| TP53 | 0.1777 | -0.085 | $<2.22 \times 10^{-16}$ |
| FAS | 0.734 | -0.0354 | $2.9 \times 10^{-8}$ |
| BAK1 | 0.00915 | -0.0044 | 0.4 |
| BAX | 0.083 | -0.0403 | $1.3 \times 10^{-9}$ |
| RPS2 | 0.302 | -0.145 | $<2.22 \times 10^{-16}$ |
| RPS15 | 0.0869 | -0.0419 | $1.1 \times 10^{-8}$ |
| RPL23A | 0.4 | -0.195 | $<2.22 \times 10^{-16}$ |
| RPL35A | 0.05 | -0.02 | $6.5 \times 10^{-5}$ |
| HIF1A | -0.054 | 0.0262 | 0.00012 |
| VEGFA | -0.339 | 0.1637 | $<2.22 \times 10^{-16}$ |
| PCBP1 | -0.051 | 0.0249 | $4.4 \times 10^{-7}$ |
| PCBP2 | 0.1003 | -0.048 | $<2.22 \times 10^{-16}$ |
| PSMB8 | 0.1349 | -0.065 | $<2.22 \times 10^{-16}$ |
| PSBM9 | 0.253 | -0.122 | $<2.22 \times 10^{-16}$ |
| TAP1 | 0.103 | -0.049 | $6.4 \times 10^{-11}$ |
| TAP2 | 0.02 | 0.009 | 0.12 |
| JAK1 | -0.060 | 0.029 | $4.5 \times 10^{-7}$ |
| JAK2 | 0.0379 | -0.0183 | 0.0072 |
| STAT1 | 0.11 | -0.056 | $1.5 \times 10^{-7}$ |
| STAT2 | 0.028 | -0.13 | 0.031 |
| MX1 | 0.1489 | -0.07 | $5.2 \times 10^{-12}$ |
| MX2 | 0.0793 | -0.038 | $2.7 \times 10^{-5}$ |
| ISG15 | 0.436 | -0.210 | $<2.22 \times 10^{-16}$ |
| IFIT3 | 0.167 | -0.08 | $2.5 \times 10^{-10}$ |
